# Cerebrovascular insulin receptors are defective in Alzheimerˈs disease

**DOI:** 10.1101/2021.12.01.470582

**Authors:** M. Leclerc, P. Bourassa, C. Tremblay, V. Caron, C. Sugère, V. Emond, D.A. Bennett, F. Calon

## Abstract

Central response to insulin is suspected to be defective in Alzheimer’s disease (AD), but its localization in the brain remains unknown. While most insulin is secreted in the bloodstream by the pancreas, how it interacts with the blood-brain barrier (BBB) to alter brain function remains poorly defined.

Here, we show that human and murine cerebral insulin receptors (INSR), particularly the long isoform INSRα-B, are concentrated in microvessels rather than in the parenchyma. Vascular concentrations of INSRα-B were lower in the parietal cortex of subjects diagnosed with AD, positively correlating with cognitive scores, leading to a shift toward a higher INSRα-A/B ratio, consistent with cerebrovascular insulin resistance in the AD brain. Vascular INSRα was inversely correlated with β-amyloid (Aβ) plaques and β-site APP cleaving enzyme 1 (BACE1), but positively correlated with insulin-degrading enzyme (IDE), neprilysin and ABCB1. Using brain cerebral intracarotid perfusion, we found that the transport rate of insulin across the BBB remained very low (<0.03 µl.g^-1^.s^-1^) and was not inhibited by an INSR antagonist. However, intracarotid perfusion of insulin induced the phosphorylation of INSRβ which was restricted to microvessels. Such an activation of vascular INSR was blunted in 3xTg-AD mice, suggesting that AD neuropathology induces insulin resistance at the level of the BBB.

Overall, the present data in postmortem AD brains and an animal model of AD indicate that defects in the INSR localized at the BBB strongly contribute to brain insulin resistance in AD, in association with Aβ pathology.

**Highlights:**

- Circulating insulin activates brain insulin receptors in microvessels.
- BBB INSR contribute to cerebral insulin resistance in AD.
- Cognitive impairment in AD is associated with a loss of cerebrovascular INSRα-B.
- Loss of isoform INSRα-B is associated with increased BACE1 activity.

**Summary:** Leclerc et al. show that circulating insulin activates cerebral insulin receptor localized on the blood-brain-barrier level (BBB), not in the parenchyma. Experiments with human brain samples and animal models provide evidence that INSR at the BBB are impaired in Alzheimer’s disease, thereby contributing to brain insulin resistance.

## Introduction

The human brain is sensitive to insulin, a hormone essential to life that is also at the heart of the pathophysiology and treatment of diabetes (Lewis and Brubaker, 2021; Polyzos and Mantzoros, 2021). The scientific literature is replete with studies in humans and other species reporting the effects of insulin on memory, cerebral blood flow, eating behavior and regulation of whole-body metabolism, supporting a therapeutic potential in brain-dependent metabolic disorders (Creo et al., 2021; Heni et al., 2015; Kleinridders et al., 2014). Although local synthesis has been detected, most insulin exercising an effect on the brain circulates in the blood after being produced by the pancreas (Banks, 2004; Csajbok and Tamas, 2016; Rhea and Banks, 2021). The blood-brain barrier (BBB) is a major interface between the blood and brain, controlling access to cerebral tissues. Therefore, to exert an effect in the central nervous system (CNS), circulating insulin must first interact with its insulin receptor (INSR) located on brain capillary endothelial cells (BCEC) forming the BBB (Banks et al., 2012; Frank and Pardridge, 1981; Gray et al., 2014; Pardridge et al., 1985; Rhea and Banks, 2021).

INSR is a disulfide-linked homodimer (αβ)2 that structurally and genetically belongs to the class II of receptor tyrosine kinases. Alternative splicing produces two isoforms of the α-chain: the short A isoform (INSRα-A) truncated by 12 amino acids (exon 11) and the long B isoform (INSRα-B) (Belfiore et al., 2017; Kellerer et al., 1992; Mosthaf et al., 1990; Yamaguchi et al., 1993). The regulation of this alternative splicing is not fully elucidated but appears to be tissue-specific (Belfiore et al., 2017; Breiner et al., 1993; Garwood et al., 2015; Malakar et al., 2016; Spencer et al., 2018). Binding of an agonist to the extracellular α-chain triggers autophosphorylation of the transmembrane β-chain, which contains multiple phosphorylation sites leading to the activation of INSR and downstream signalization pathways (Lee and Pilch, 1994; Tornqvist et al., 1987; Weiss and Lawrence, 2018).

Beside classical amyloid-β (Aβ) and tau pathologies, Alzheimer’s disease (AD) is characterized by defective brain uptake of glucose and impaired response to insulin (Arnold et al., 2018; Baglietto-Vargas et al., 2016; de la Monte, 2019; Hoyer, 2004; Kellar and Craft, 2020; Stanley et al., 2016). The most compelling evidence come from postmortem indexes of brain insulin resistance, such as changes in INSR or IRS1 phosphorylation status shown to be associated with cognitive impairment in humans and in mice (Arnold et al., 2014; Arvanitakis et al., 2020; Denver et al., 2018; Talbot et al., 2012). Insulin administration in animals is reported to improve memory function, including in mouse models of AD (Mao et al., 2016; Sanguinetti et al., 2019; Vandal et al., 2014). Using intranasal delivery to avoid hypoglycemic effect of systemically injected insulin, several small clinical trials have reported benefits on memory scores in cognitively impaired adults (Claxton et al., 2015; Craft et al., 2017; Reger et al., 2008). However, recent larger clinical trials brought mitigated results (Craft et al., 2020; Gwizdala et al., 2021; Kellar et al., 2021; Rosenbloom et al., 2021). Still, increasing insulin sensitivity remains a promising avenue and a wide range of repurposed diabetes drugs are in ongoing clinical trials, including metformin, thiazolidinediones and glucagon-like peptide 1 (GLP-1) analogs (Cummings et al., 2020; Hölscher, 2021; Moran et al., 2019).

While cerebral insulin resistance is often assumed to be restricted to neurons in AD (Frazier et al., 2020; Lourenco et al., 2015; Moloney et al., 2010), it is supported by limited immunohistochemical (IHC) evidence (Moss et al., 1990; Unger et al., 1989; Unger et al., 1991). Most initial studies on the distribution of brain INSR used macroscopic techniques (Baskin et al., 1986; Dore et al., 1997; Frolich et al., 1998; Kar et al., 1993; Marks et al., 1990). However, recent single-cell transcriptomic analyses indicate that the mRNA transcript encoded by the *INSR* gene is found in higher concentrations in endothelial cells in the mouse and human brain (Vanlandewijck et al., 2018; Yang et al., 2021; Zhang et al., 2020). Accordingly, a growing number of studies are showing that INSR located in the cerebral vasculature plays a key role in the action of insulin on the brain (Konishi et al., 2017; Rhea and Banks, 2019; Vicent et al., 2003). Based on the current understanding, INSR at the BBB binds circulating insulin to either (i) act as a classic receptor to trigger cell-signaling pathways within BCEC or (ii) as a transporter to ferry an insulin molecule into the brain parenchyma (Banks and Kastin, 1998; Baura et al., 1993; Duffy and Pardridge, 1987; Gray et al., 2017; Woods et al., 2003). So far, it remains unclear whether the same INSR protein can exert both roles (Gray and Barrett, 2018; Hersom et al., 2018; Rhea and Banks, 2021; Rhea et al., 2018).

Given this large set of clinical and preclinical data showing a primary role of insulin on brain function, we aimed to determine how circulating insulin interacts with the BBB INSR and whether the latter is defective in AD. We first sought to determine INSR levels in microvascular extracts from both mouse and human brain samples relative to brain parenchyma. To probe for changes in AD, we used microvessel extracts of parietal cortex from participants in the Religious Orders Study who underwent detailed clinical and neuropsychological evaluations (Bennett et al., 2018; Bourassa et al., 2019b; Bourassa et al., 2020). Association with clinical and biochemical data were assessed, including BBB transporters and receptors putatively involved in Aβ transport. To directly investigate the response of brain INSR to circulating insulin, we performed intracarotid insulin perfusion and found that INSR activation is localized at the BBB, where it is impaired by Aβ and tau pathologies in a mouse model of AD at different ages.

## Materials and methods

### Human samples: Religious Orders Study (ROS) (Rush Alzheimer’s Disease Center)

Parietal cortex samples were obtained from participants in the ROS, a longitudinal clinical and pathological study of aging and dementia (Bennett et al., 2018; Bourassa et al., 2019b; Bourassa et al., 2020). Each participant enrolled without known dementia and agreed to an annual detailed clinical evaluation and brain donation at death. The study was approved by an Institutional Review Board of Rush University Medical Center. All participants signed an informed consent, an Anatomic Gift Act for brain donation, and a repository consent allowing their data and biospecimens to be shared. A total of 21 cognitive performance tests were administered of which 19 were used to create a global measure of cognition, and five cognitive domains: episodic, semantic, working memory, perceptual speed and visuospatial ability (Bennett et al., 2006a; Wilson et al., 2002). Participants received a clinical diagnosis of Alzheimer’ dementia (AD), mild cognitive impairment (MCI) or no cognitive impairment (NCI) (n = 20 for each group) at the time of death by a neurologist, blinded to all postmortem data, as previously described (Bennett et al., 2006a; Bennett et al., 2006b; Bennett et al., 2002). In addition, current prescription medication usage in the last two weeks prior to each evaluation were available, such as antihypertensive and diabetes medications (Arvanitakis et al., 2008a; Arvanitakis et al., 2008b).

The neuropathological diagnosis is based on the ABC scoring method found in the revised National Institute of Aging – Alzheimer’s Association (NIA-AA) guidelines for the neuropathological diagnosis of AD (Montine et al., 2012). Four levels of AD neuropathological changes (not, low, intermediate, or high) were determined using (A) Thal score assessing phases of Aβ plaque accumulation (Thal et al., 2002), (B) Braak score assessing neurofibrillary tangle pathology (Braak and Braak, 1991), and (C) CERAD score assessing neuritic plaque pathology (Mirra et al., 1991). Individuals classified as “Controls“ had no or a low level of AD neuropathological changes while those classified as “AD“ included participants with intermediate or high levels of AD neuropathological changes (Bourassa et al., 2019b). Neuritic plaques and neurofibrillary tangles in the parietal cortex were counted following Bielschowsky silver impregnation, as previously described (Bennett et al., 2003). As previously described (Bourassa et al., 2019b; Tremblay et al., 2017; Tremblay et al., 2007), soluble and insoluble levels of Aβ40 and Aβ42 were also assessed in homogenates and microvascular extracts of parietal cortex. See **Suppl. Table 1** for the characteristics of the cohort.

### Isolation of brain microvessels

Human brain microvessels were extracted following the procedure described previously (Bourassa et al., 2019b). Following a series of centrifugation steps, including a density gradient centrifugation with dextran and a filtration (20-µm nylon filter), two fractions were obtained: one enriched in cerebral microvessels, the other consisting of microvessel-depleted parenchymal cell populations. Examples of the efficiency of the separation are shown in **Fig. 1**, using immunodetection of endothelial and neuronal markers (Bourassa et al., 2019b). Proteins of both fractions were extracted using a lysis buffer (150 mM NaCl, 10 mM NaH2PO4, 1 mM EDTA, 1% Triton X-100, 0.5% SDS and 0.5% deoxycholate) containing the same protease and phosphatase inhibitors, and supernatants were kept for Western immunoblotting analyses. Brain microvessels from mice were generated with a similar procedure reported in previous work (Bourassa et al., 2019b; Bourassa et al., 2020; Traversy et al., 2017). A scheme is shown in **Suppl. Fig. 1.**

**Figure 1.**
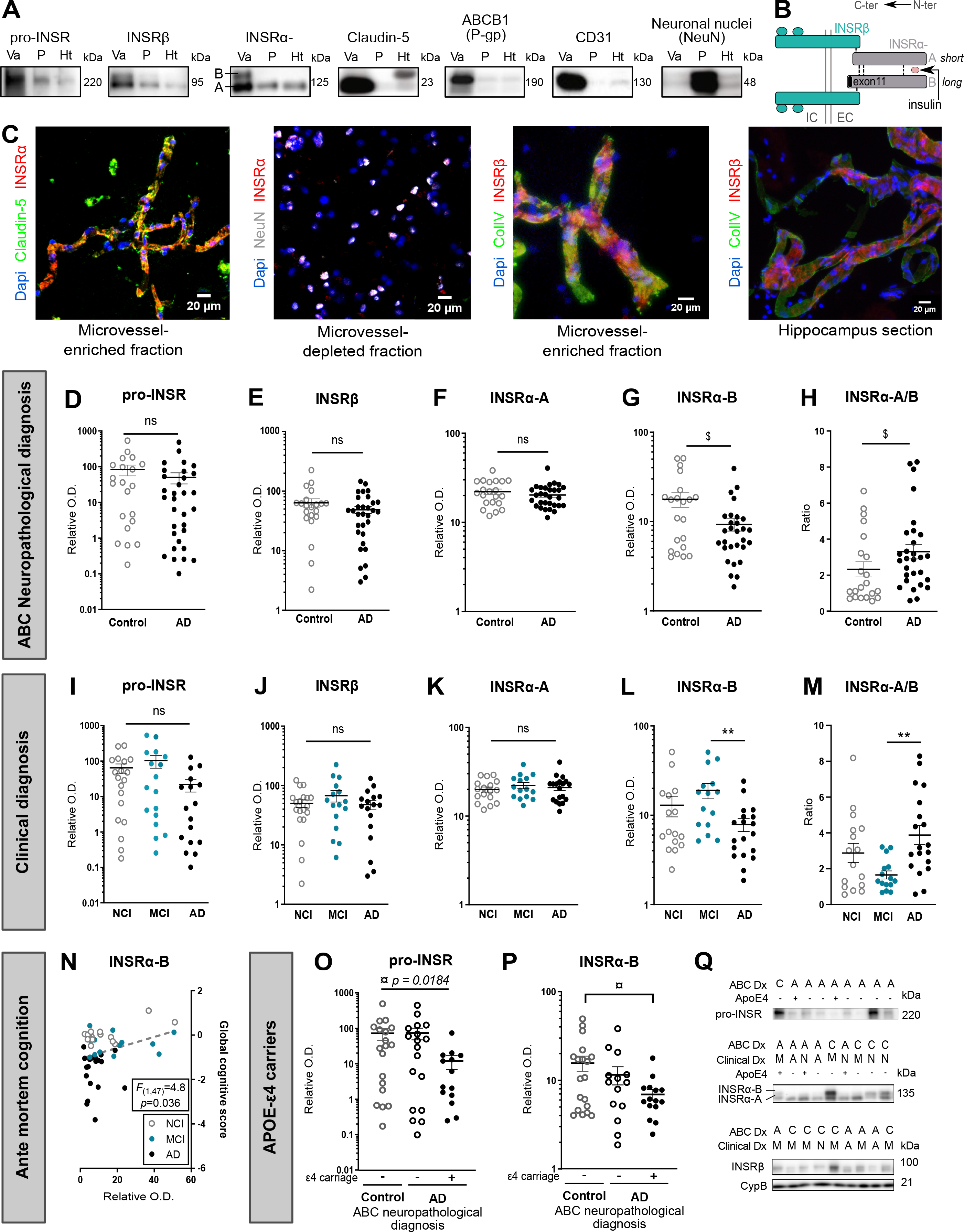
INSRα-B levels are reduced in Alzheimer’s disease, correlating with cognitive dysfunction. (A) Pro-INSR, INSRα and INSRβ are enriched in vascular fractions from the parietal cortex, along with endothelial markers claudin-5, ABCB1(P-gp) and CD31, whereas Neuronal nuclei (NeuN), a neuronal marker, is rather concentrated in the microvessel-depleted parenchymal fraction. (B) Schematic representation of the transmembrane INSR: the extracellular α chain binds circulating insulin and the intracellular β chain acts as a primary effector of insulin signaling pathway through auto- phosphorylation. INSRα is expressed as isoforms A and B due to alternative splicing (exon 11 is spliced for isoform B). (C) Human INSRα is colocalized with claudin-5 and collagen IV in brain microvessels and hippocampal section, whereas no colocalization was observed with NeuN-labeled neurons in microvessel-depleted fractions (Magnification used: 20×, scales bar: 20 μm). (D-M) Dot plots of the concentrations of INSR forms in microvessels comparing participants based on neuropathological diagnosis following the ABC criteria (D-H) or clinical diagnosis (I-M). Unpaired t-test ($p < 0.05) or one-way ANOVA followed by Tukey’s post-hoc (**p < 0.01). (N) Correlation between the levels of vascular INSRα-B and the global cognitive score. Linear regression analyses were controlled for educational level, age at death, sex and apoE genotype. *F* ratio and p-value are shown. (O,P) Dot plots of the concentrations of pro-INSR and INSRα-B in microvessels comparing APOE4 carriers based on the ABC neuropathological diagnosis. Welch-ANOVA followed by Dunnett’s post-hoc (¤p<0.05). (Q) Representative Western blots of consecutive bands are shown. Data were log transformed for statistical analysis and are represented as scatter plots with a logarithmic scale. Horizontal bars indicate mean ± SEM. Abbreviations: ABC: Dx Neuropathological Diagnosis, ABCB1(P-gp): ATP Binding Cassette Subfamily B Member 1/P-glycoprotein, A-AD: Alzheimer’s disease, ApoE: Apolipoprotein E genotype, ColIV: Collagen- IV, C: Control, CD31: Platelet endothelial cell adhesion molecule, Clinical Dx: Clinical Diagnosis, CypB: Cyclophilin B, EC: Extracellular, Ht: Total homogenate, IC: Intracellular, INSR: Insulin receptor, M-MCI: Mild cognitive impairment, NeuN: Neuronal nuclei, N-NCI: Healthy controls with no cognitive impairment, O.D.: Optical density, P: Microvessel-depleted parenchymal fraction, Va: Vascular fraction enriched in microvessels.

### Immunofluorescence analysis of isolated human microvessels

The method was similar to previous publications (Bourassa et al., 2019b; Bourassa et al., 2020). Both vascular and microvessel-depleted extracts on glass side were fixed using a 4% paraformaldehyde solution in phosphate buffer saline (PBS) for 20 minutes at room temperature (RT) and then blocked with a 10% normal horse serum and 0.1% Triton X-100 solution in PBS or commercial TrueBlack® Background Suppressor and Blocking Buffer (Biotum) for 1 h at RT. For Aβ immunolabeling, an additional pretreatment with 90% formic acid during 10 min was performed between the fixation and blocking steps. Following an incubation overnight at 4 °C with primary antibodies diluted in a 1% NHS and 0.05% Triton X-100 solution in PBS, or TrueBlack® Blocking Buffer (Biotum), vascular and microvessel-depleted extracts were incubated with secondary antibodies diluted in the same solution as the primary antibodies during 1 h at RT. Primary and secondary antibodies are listed in **Suppl. Table 2**. Cell nuclei were counterstained with DAPI (Thermo Fisher Scientific, 0.02% in PBS) and slides were mounted with Mowiol mounting medium or Prolong Diamond antifade (Thermo Fisher Scientific). If needed, slides were incubated with TrueBlack Plus (Biotum) and TrueView (Biotum) to decrease autofluorescence and background. Between each step, three washes of 5 min in PBS were performed.

Images were taken using a fluorescence microscope (EVOS fl Autoimaging system; Thermo Fisher Scientific) at magnification 20X-40X or a laser scanning confocal microscope (Olympus IX81, FV1000; Ontario, Canada) with sequential scanning acquisition using optimal z-separation at a magnification of 20X.

### Animals

The use of animals was approved by the Laval University animal research committee (CPAUL) in accordance with the standards of the Canadian Council on Animal Care. All animals had free access to laboratory food and water and were maintained under a 12-h light–dark cycle at 22°C.

The 3xTg-AD mouse is a widely used model that progressively develops both Aβ deposits and neurofibrillary tangles (τ pathology) as well as cognitive deficits, due to the expression three mutant genes: β-amyloid precursors protein (APPswe), presenilin-1 (PS1MA146V) and tau (tauP301L) (Julien et al., 2010; Oddo et al., 2003; Phivilay et al., 2009). It fully develops neuropathological and behavioral changes around 12 months of age (Arsenault et al., 2013; Belfiore et al., 2019; Dal-Pan et al., 2017; Vandal et al., 2015; Vandal et al., 2016). Previous studies showed that 3xTg-AD mice also display metabolic disorders (Bosoi et al., 2021; Julien et al., 2010; Nicholson et al., 2010; Sanguinetti et al., 2019; Vandal et al., 2015; Vandal et al., 2014; Velazquez et al., 2017), and their brain Aβ concentrations quickly decrease after insulin administration (Chen et al., 2014; Vandal et al., 2014). Here, they were compared with non transgenic (Non-Tg) littermates with the same genetic background (C57Bl6/129SvJ). All groups of 3xTg-AD and Non-Tg mice were composed of males and females in similar proportion (42 males and 34 females were used for comparison of microvessel proteins between 6, 12 18 months of age and 35 males and 30 females for intracarotid *in situ* cerebral perfusion (ISCP)). In addition, 6-month-old C57Bl6 mice (n=24 with 11 males and 13 females) were used for insulin stimulation experiments, using ISCP followed by microvessel extraction (see **Suppl. Fig. 1**). In addition, 15-weeks-old Balb/c (N=22) or C57Bl/6 mice (N=7) for insulin transport quantification by ISCP.

Animals were sacrificed by intracardiac perfusion or ISCP (for ISCP, see below). Intracardiac perfusion was performed with phosphate saline buffer containing a cocktail of protease inhibitors (SIGMAFAST^TM^ Protease Inhibitor Tablets, Sigma-Aldrich, St Louis, MO, USA) and phosphatase inhibitors (1 mM sodium pyrophosphate and 50 mM sodium fluoride) under deep anesthesia with ketamine/xylazine i.p. (300/30 mg/kg).

### Acute exposition of the BBB to insulin using intracarotid *in situ* cerebral perfusion (ISCP)

The intracarotid ISCP technique consists in a direct cerebral infusion of a labeled compound at the blood-brain barrier through the right common carotid artery (Alata et al., 2014; Bourassa et al., 2019a; Do et al., 2014). Briefly, mice were anesthetized by intraperitoneal injection of a mixture of ketamine/xylazine i.p. (140/8 mg/kg). The right external and the right common carotid artery were ligated, and the common carotid artery was catheterized with polyethylene tubing (0.30 mm x 0.70 mm) filled with heparin (25 IU/ml). The syringe containing the perfusion fluid was placed in an infusion pump and connected to the catheter. The thorax was opened, the heart cut and perfusion started immediately. The perfusion fluid contained bicarbonate buffered physiologic saline containing 128 mM NaCl, 24 mM NaHCO3, 4.2 mM KCl, 2.4 mM NaH2PO4, 1.5 mM CaCl2, 0.9 mM MgCl2, and 9 mM D-glucose. The solution was gassed with 95% O2–5% CO2 for pH control (7.4), filtrated (0.20µm) and used at 37°C. For radiolabeled insulin transport experiment mice were perfused with 0.1 nM of ^125^I-insulin, 0.3 µCi/ml of ^3^H-sucrose as a vascular marker, and 20 nM S961 for competition. The physiological buffer was supplemented with 0.25% bovine serum albumin (BSA) and the perfusion lasted 120 sec at 1.25 ml/min. For insulin stimulation studies, 6-month-old C57Bl6 and 16-month-old Non-Tg and 3xTg-AD mice were perfused for 120 sec, at 2.5 ml/min, with insulin (100 or 350 nM) or saline.

### Western blot

For Western immunoblotting, proteins from the vascular fraction were extracted and a protein quantification was done using bicinchoninic acid assays (Pierce, Rockford, IL). Protein homogenates from human and murine brain microvascular extracts were added to Laemmli’s loading buffer and samples where heated 10 minutes at 70°C (human and 3xTg-AD samples) or not (samples from insulin stimulation studies). Equal amounts of proteins per sample (4 to 8 µg) were resolved on a sodium dodecyl sulfate-polyacrylamide gel electrophoresis (SDS-PAGE) 8% acrylamide. All samples, loaded in a random order, were run on the same immunoblot experiment for quantification.

Proteins were electroblotted on PVDF membranes, which were then blocked during 1h at RT with Superblock™ in PBS blocking buffer (Thermo fisher Scientific #37515) or PBS containing 0.1% Tween 20 and 3% BSA. If needed, membranes were stained with No-Stain™ protein labeling reagent (Thermo Fisher scientific #A44717) to assess a similar protein load in the samples. Membranes were then incubated overnight at 4°C with primary antibodies (See **Suppl. Table 2**). Finding satisfactory primary antibodies for INSR epitopes and downstream signaling elements, particularly in human postmortem samples, was challenging and those that failed to give conclusive results are listed in **Suppl. Table 3**. Membranes were then washed three times with PBS containing 0.1% Tween 20 and incubated during 1h at RT with the secondary antibody in PBS containing 0.1% Tween 20 and 1% BSA. Membranes were probed with chemiluminescence reagent (Luminata Forte Western HRP substrate; Millipore #ELLUF0100) and imaged using the myECL imager system (Thermo Fisher Scientific) or the Amersham Imager 680 (GE Healthcare Bio-Sciences). Densitometric analysis was performed using the Image Lab™ Software (Biorad). Uncropped gels of immunoblotting experiments conducted with human samples are shown in **Suppl. Fig.2**.

### ELISA for human Aβ and APP-βCTF

Concentrations of β-secretase-derived βAPP fragment (APP-βCTF or C99) in detergent-soluble extracts of brain microvessels (parietal cortex) were determined using Human APP-βCTF ELISA Assay kit from IBL (Gunma, Japan) according to the manufacturer’s instructions.

### Data and statistical analysis

Statistical analyses were performed using Prism 9.0 (GraphPad Software Inc.) and JMP 15 (SAS) software. The threshold for statistical significance was set to p < 0.05. Homogeneity of variance and normality were determined for all data sets using D’Agostino & Pearson’s normality test or Shapiro-Wilk test. When normality was verified, unpaired Student’s t-tests were used to identify significant differences between two groups. Otherwise, Welch correction or a Mann-Whitney tests were performed. When more than two groups were compared, parametric one-way ANOVA followed by Tukey’s multiple comparison tests were performed, unless variances were different, in which case Welch-ANOVA followed by Dunnett’s multiple comparison tests were used. When both normality and variance equality were not confirmed, a non- parametric Kruskall-Wallis test followed by Dunn’s multiple comparison was performed. When needed, logarithmic transformation was performed to reduce variances and provide more normally distributed data. Multivariate analyses were employed to assess association between continuous variables. Significance of correlations with antemortem clinical scores were adjusted for the following covariates: education level, sex, age at death, and Apolipoprotein E (APOE) genotype.

### Data availability

The data supporting the findings of this study are available from the corresponding author upon reasonable request. Data from the Religious Orders Study can be requested at https://www.radc.rush.edu.

## Results

### INSRs are enriched in human brain microvessels: levels of the isoform INSRα-B are reduced in AD

To determine the postmortem localization of the INSR in the human brain, we first extracted microvessels from parietal cortex samples collected from ROS participants. We opted for a soft procedure using sequential centrifugation and filtration steps, adapted for frozen samples, as previously described (Bourassa et al., 2019b; Bourassa et al., 2020). We observed that INSR were present in higher concentrations in vascular fractions (Va) containing brain microvessels, compared to microvessel-depleted parenchymal fractions (P) and whole brain homogenates (Ht) (**Fig. 1A**). The validity of the method was confirmed by the coinciding enrichment in endothelial markers claudin-5, ABCB1 (P-gp) and CD31 in the same vascular fractions, whereas neuronal nuclei (NeuN) staining was more prominent in microvessel-depleted fractions (**Fig. 1A**), as shown in our previous work (Bourassa et al., 2019b; Bourassa et al., 2020).

INSR is a heterotetrametric receptor combining two dimers composed of both α (extracellular) and β (intracellular) chains, recognizable by different antibodies (**Fig. 1B and Suppl. Tables 2, 3**) but all coded from the same *INSR* gene. INSRα incorporates the binding site for circulating insulin and exists in two isoforms depending on the absence or presence of exon 11, depicted as A and B, respectively. The full-size precursor of the INSR, pro-INSR, and both INSRα and INSRβ isomers were enriched in vascular fractions (**Fig. 1A**).

The localization of INSRα on microvessels was confirmed by confocal immunofluorescence where the signal of human INSRα antibody overlapped with claudin-5, a tight-junction protein of brain endothelial cells, but was almost absent in microvessel-depleted fractions containing NeuN-positive cells. In addition, immunosignals for INSRβ and collagen IV, a basal membrane protein, colocalized in human microvessels from microvascular-enriched fractions and brain sections (**Fig. 1C**).

Western blot analyses revealed that INSRα-B levels were lower in vascular fractions from participants with a neuropathological (-48%) or clinical (-40% and -59% compared to C and MCI, respectively) diagnosis of AD (**Fig. 1G, L**), whereas levels of pro-INSR, INSRβ or INSRα-A did not differ (**Fig. 1D-F, I-K**). A lower INSRα- B was also found in association with Thal, Braak and CERAD ratings (**Suppl. Fig. 3**). In the parenchymal fractions, where the INSRα B-isoform was absent, levels of the INSRα A-isoform were not different between groups (**Suppl. Fig. 4**). Interestingly, persons with a neuropathological or clinical diagnosis of AD had a higher vascular INSRα-A/B ratio (**Fig. 1H, M**), a molecular index of insulin resistance in the liver and adipose tissue (Belfiore et al., 2017; Besic et al., 2015; Chettouh et al., 2013; Escribano et al., 2009; Kaminska et al., 2014). In addition, *ante mortem* global cognition was positively correlated to the cerebrovascular content in INSRα-B (**Fig. 1N**), but not to other INSR isoforms (**Suppl. Fig. 5**). Finally, AD subjects carrying one apoE4 allele had lower concentrations of pro-INSR (-83% and -83%) and INSRα-B (- 40% and -56%) compared to AD non-carriers and control subjects respectively (**Fig. 1O, P**). Immunoblots are shown in **Fig. 1Q** and in **Suppl. Fig. 2**.

### Levels of vascular INSR correlate with Aβ pathology or BBB proteins involved in Aβ production and clearance

We next investigated the association between vascular INSR and several markers of Aβ pathology previously assessed in whole homogenates and/or in microvascular extracts from the parietal cortex from the same series of ROS participants (**Fig. 2A**)(Bourassa et al., 2019b; Tremblay et al., 2017; Tremblay et al., 2007). Significant inverse correlations were found between vascular levels of INSRα-B, neuritic plaques, as well as vascular β-secretase (BACE1), an enzyme implicated in APP cleavage and Aβ formation (**Fig. 2B, C**). By contrast, INSRα-B levels in vascular extracts were positively correlated with vascular neprilysin and insulin degrading enzyme (IDE), two enzymes involved in Aβ degradation (**Fig. 2A, D**). In addition, strong associations were detected between vascular INSRα-B, ABCB1 (P-gp) and LRP1, implicated in Aβ efflux (Deane et al., 2004; Storck et al., 2016), suggesting that more INSRα-B at the BBB is associated with a higher clearance of Aβ from brain to blood (**Fig. 2A, E**). However, soluble and insoluble levels of Aβ40 or Aβ42 in parietal cortex proteins homogenates did not show any association with vascular INSRα-B. No association was found with all soluble or insoluble markers of tau pathology tested, except a weak inverse correlation between vascular INSRβ and neurofibrillary tangles (**Fig. 2A**). Finally, we noted than INSRα-B and most of the other INSR isoforms were positively associated with caveolin-1 and eNOS, two proteins involved in receptor recycling and endothelial generation of nitric oxide, respectively (**Fig. 2A, F, G**).

**Figure 2.**
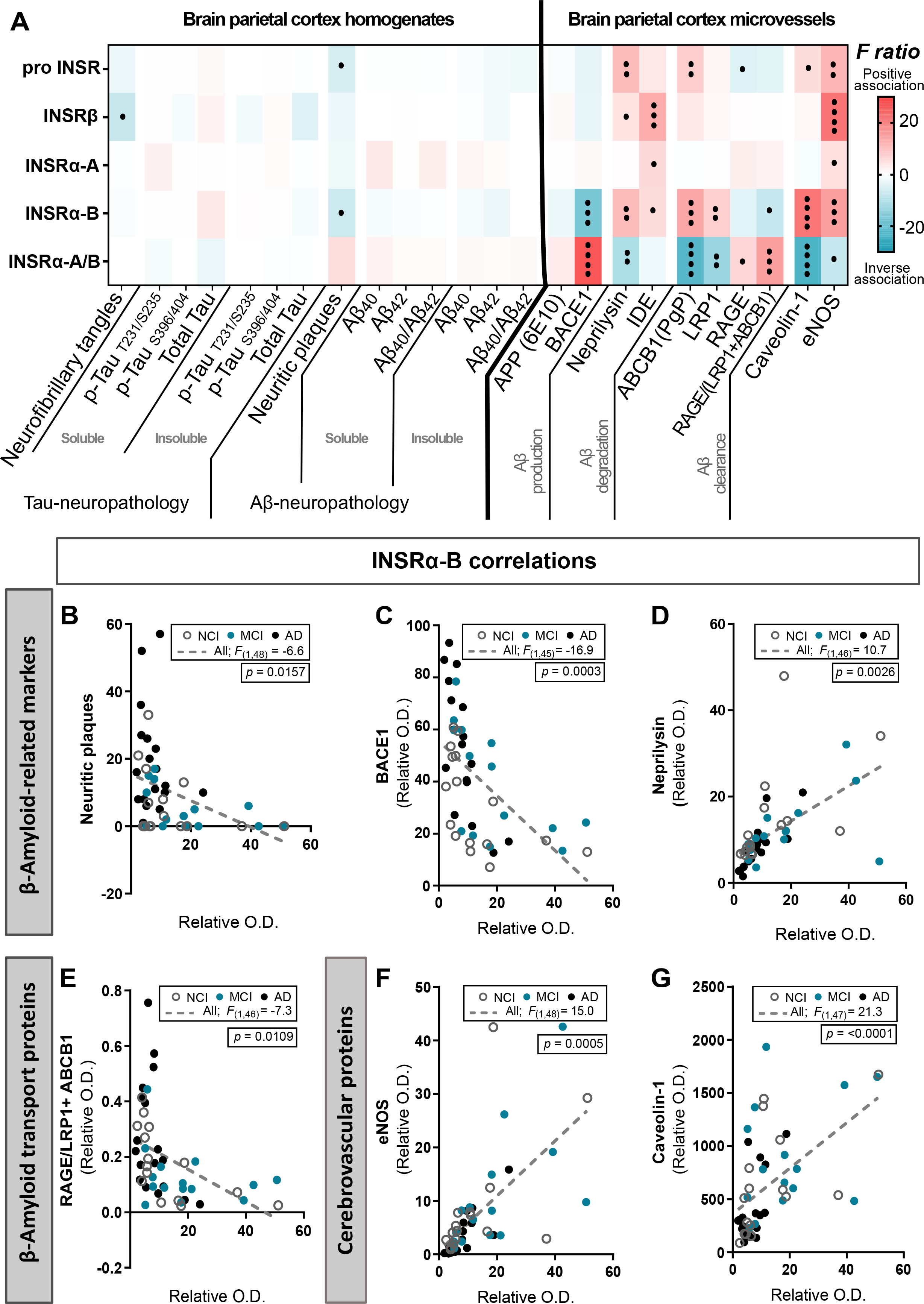
Cerebrovascular levels of INSRα-B correlate with neuritic plaques and Aβ-related proteins located on the BBB. (A-G) Correlations between cerebrovascular INSRs and neurofibrillary tangles count, phosphorylated-tau T231/S235 (AT180), S396/404 (PHF1, confirmed with AD2) (Tau-neuropathology), cortex neuritic plaques count, and Aβ concentrations (Aβ-neuropathology) in brain homogenates. Correlations were also established with other proteins assessed in microvessel-enriched fractions. Soluble and insoluble proteins are found in TBS-soluble and formic acid-soluble fractions, respectively. Linear regressions analyses were controlled for educational level, age at death, sex and ApoE genotype, and were performed to generate F ratios and p-values in the heatmap (•p < 0.05; ••p < 0.01; •••p < 0.001; ••••p < 0.0001). Red and blue highlighted cells respectively indicate significant positive and negative correlations (A). Graphical representation of noteworthy correlations between INSRα-B and Aβ-related markers (B-D) and transporters (E), as well as microvessel markers (F,G). Abbreviations: Aβ: β-amyloid protein, ABCB1(P-gp): ATP Binding Cassette Subfamily B Member 1/P- glycoprotein, APP: Amyloid Precursor Protein, BACE1: β-site APP cleaving enzyme, eNOS: Endothelial NOS, IDE: Insulin degrading enzyme, INSR: Insulin receptor, LRP1: Low density lipoprotein receptor-related protein 1, MCI: Mild cognitive impairment, NCI: Healthy controls with no cognitive impairment, O.D.: Optical density, RAGE: Receptor for Advanced Glycation End products.

### INSR is altered in 3xTg-AD mouse microvessels

To assess the localization of INSR in the mouse brain, we performed the same extraction procedures used with human samples. We observed a vascular enrichment for all isoforms, but stronger for pro-INSR and INSRα-B, compared to total homogenates or microvessel-depleted parenchymal fractions (**Fig. 3A**), corroborating human data. The concomitant enrichment of endothelial markers claudin-5, CD31 and ABCB1 (P-gp) in vascular fractions validated the method.

**Figure 3.**
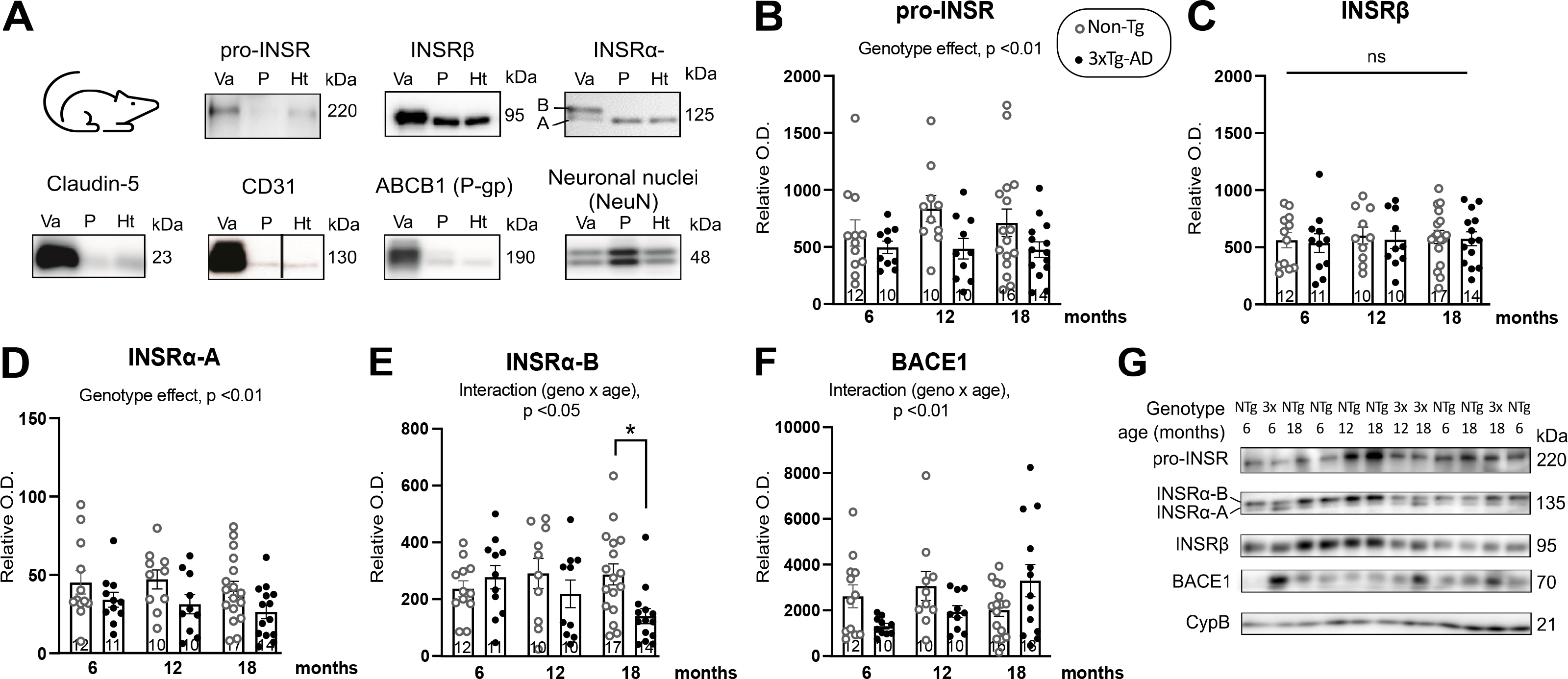
Lower vascular INSR in old 3xTg-AD mice. (A) Representative Western blots showing enriched content in pro-INSR, INSRα and INSRβ in vascular fractions compared to parenchymal fractions (rich in neuronal marker NeuN) and total homogenates from the mouse brain, along with known endothelial markers claudin-5, CD31 and ABCB1(P-gp). (B-F) Vascular levels of pro-INSR, INSRβ, INSRα-A, INSRα-B and BACE1 in 3xTg-AD and Non-Tg mice at 6, 12 and 18 months of age (42 males and 34 females). Two-way ANOVA followed by Tukey’s post-hoc test (*p < 0.05; ns, non-significant), and linear trend model. Data are presented as mean ± SEM. (G) Representative Western blots of consecutive bands are shown Abbreviations: 3x-3xTg-AD: tri-transgenic mice, ABCB1(P-gp): ATP Binding Cassette Subfamily B Member 1/P-glycoprotein, BACE1: β-site APP cleaving enzyme, CD31: Platelet endothelial cell adhesion molecule, CypB: Cyclophilin B, EC: Extracellular, Ht: Total homogenate, IC: Intracellular, INSR: Insulin receptor, NeuN: Neuronal nuclei, NTg-Non-Tg: Non-transgenic mice, O.D.: Optical density, P: Microvessel-depleted parenchymal fraction, Va: Vascular fraction enriched in microvessels.

To determine whether changes in INSR are a consequence of classical AD neuropathology, we applied the same methodology to brain samples from 3xTg-AD mice at different ages (6, 12 and 18 months). Multivariate analyses showed that the 3xTg-AD genotype was associated with lower INSRα-A or pro-INSR levels, although difference at each age remained non-significant (**Fig. 3B, D, G**). Similar to the findings in human microvessel extracts, no difference between groups was observed for INSRβ (**Fig. 3C, G**). Interestingly, INSRα-B levels in microvessel-enriched fractions were lower in 3xTg-AD mice at 18 months of age compared to non-transgenic mice (-51.1%), in accordance with human data (**Fig. 3E, G**). A significant linear trend from 6 to 18 months (r^2^ =-0.1828, p=0.0116) and a genotype-age interaction were detected, consistent with an age-induced decrease of INSRα-B only in 3xTg-AD animals (**Fig. 3E, G**). By contrast, we also found a significant genotype-age interaction for BACE1, associated with a linear upward trend with age of vascular BACE1 levels in 3xTg-AD mice (r^2^ =0.2130, p=0.0076)(**Fig. 3F, G**). These results unveil a pattern of changes of INSR, induced by age and Aβ/tau pathologies, in agreement with human data.

### The insulin-induced activation of INSR at the BBB is blunted in 3xTg-AD mice

Our results so far indicate that cerebral INSR are concentrated in microvessels, but with levels decreasing along with AD pathology. Previous published data suggest that AD is associated with reduced activation of INSR and its downstream intracellular signaling (Arnold et al., 2018; Arvanitakis et al., 2020; Talbot et al., 2012). However, it remains to be shown where blood-borne insulin activates INSRs in the brain, and whether this activation is disturbed by AD pathology. Like other members of the tyrosine kinase family of receptors, the binding of an agonist triggers autophosphorylation of the INSR. However, its implication as a transporter to ferry circulating insulin into the brain parenchyma has been recently disputed (Rhea and Banks, 2021). We thus directly assessed the transport of insulin through the BBB using intracarotid ISCP, with or without an INSR-specific competitive antagonist S961 (Knudsen et al., 2012; Schaffer et al., 2008)(**Fig. 4A**). Perfused ^125^I-insulin was set at 0.1 nM, close to the physiological concentration of circulating insulin in the mouse (Ahrén et al., 1997; Flatt and Bailey, 1981). We found that the measured brain uptake coefficient of ^125^I-insulin was very low (0.028 ± 0.007 µl.g^-1^.s^-1^) compared to a highly diffusible compound such as diazepam (45.6 ± 2.6 µl.g^-1^.s^-1^). When ^125^I-insulin was perfused within a human serum, the brain uptake coefficient became negligible (0.011 ± 0.013 µl.g^-1^.s^-1^, n = 6), indicating that insulin interaction with blood component ever further decreases its brain uptake in physiological conditions. In addition, S961 (20 nM) did not block ^125^I-insulin transport. We also confirmed that neither insulin nor S961 had an impact on BBB permeability using ^3^H-sucrose (**Fig. 4B, C**). This data suggests that INSR at the BBB does not act as the main transporter for insulin, in accordance with published data evidence *in vitro* and *in vivo* (Hersom et al., 2018; Rhea et al., 2018; Williams et al., 2018).

**Figure 4.**
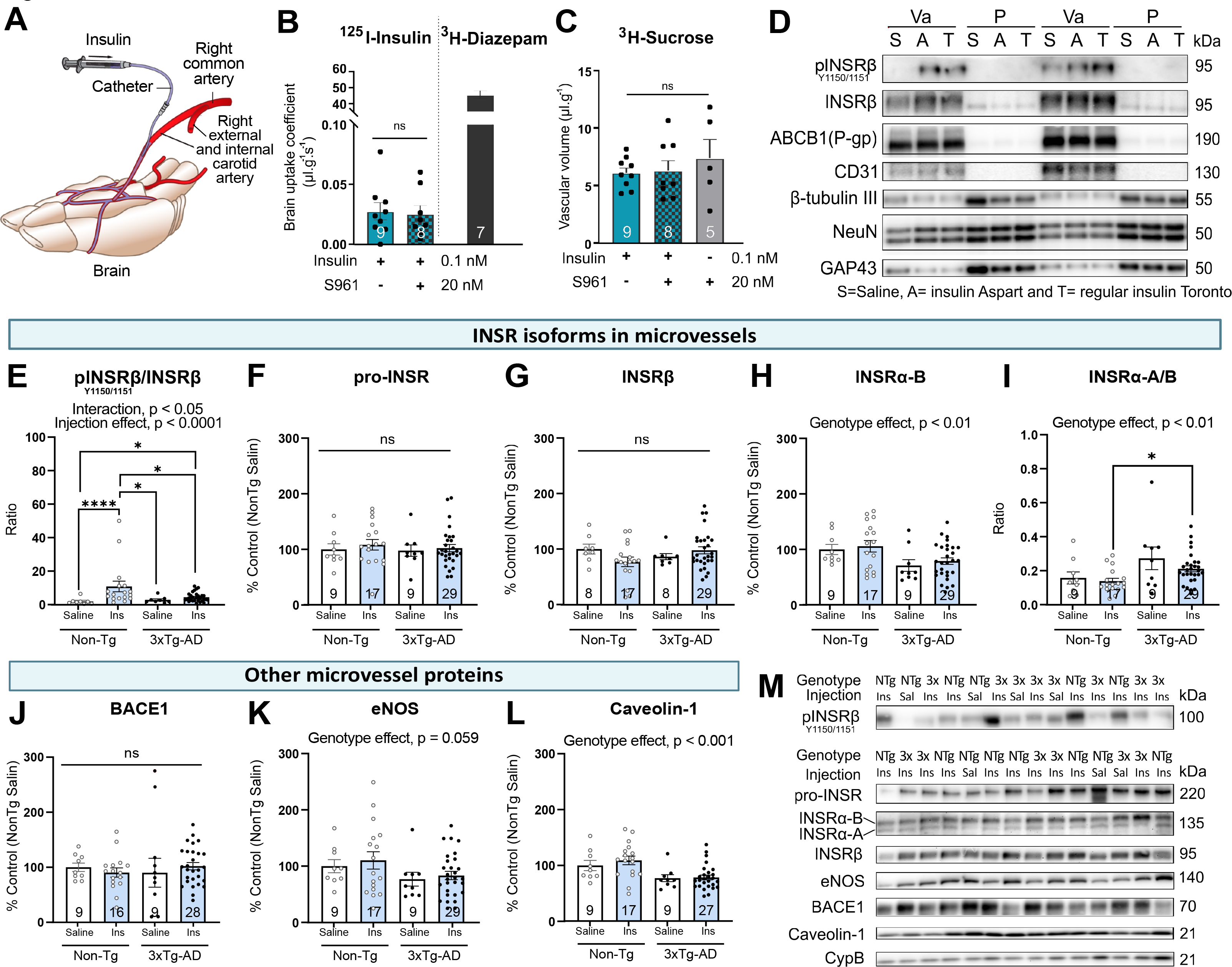
The activation of INSR after intracarotid injection of insulin is restricted to brain microvessels and is blunted in 3xTg-AD mice. (A-C) Illustration of the in situ cerebral perfusion (ISCP) technique (A). Brain uptake coefficient of ^125^I- insulin (0.01 nM) perfused alone or with S961 (20 nM), a selective INSR antagonist competing with insulin, in Balb/c mice aged 15 weeks. ^3^H-Diazepam (0.3 µCi/µl) was perfused in age-matched C57Bl6 mice to emphasize the difference between a highly diffusible drug and insulin (B). No significant change in cerebrovascular volume due to the treatment was observed by coperfusing the vascular marker ^3^H-sucrose (0.3 µCi/ml) (C). Unpaired t-test or parametric one-way analysis of variance (ns, non-significant). Data are presented as mean ± SEM. (D) Representative Western blots showing phosphorylated INSRβ in vascular fractions following perfusion of insulin (<u>A</u>spart or <u>T</u>oronto) by ISCP in 6-month-old C57Bl6 mice, compared to parenchymal fractions and total homogenates. Endothelial markers CD31 and ABCB1 (P-gp) as well as the neuronal markers β-tubulin III, NeuN and GAP43 and were immunoblotted on the same membranes. (E-L) Dot plots of the concentrations of INSR isoforms in microvessels comparing mice based on genotype and insulin perfusion (E-I), and cerebrovascular proteins BACE1, eNOS and caveolin-1 (J,L). One-way or two-way analysis of variance followed by Tukey’s post-hoc (*p<0.05;****p<0.0001). Data were log transformed for statistical analysis. Horizontal bars indicate mean ± SEM. Representative Western blots of consecutive bands are shown, from the same samples, but from a different randomization (M). Abbreviations: 3x-3xTg-AD: tri-transgenic mice, A: insulin Aspart (NovoRapid®), ABCB1(P-gp): ATP Binding Cassette Subfamily B Member 1/P-glycoprotein, BACE1: β-site APP cleaving enzyme, CD31: Platelet endothelial cell adhesion molecule, CypB: Cyclophilin B, eNOS: Endothelial NOS, Ins: insulin perfusion, GAP43: Growth Associated Protein 43, INSR: Insulin receptor, NeuN: Neuronal nuclei, NTg-Non-Tg: Non- transgenic mice, O.D.: Optical density, P: Microvessel-depleted parenchymal fraction, S961: INSR antagonist, S-Sal: saline perfusion, T: regular insulin Toronto (Novolin ge®), Va: Vascular fraction enriched in microvessels.

Then, we used ISCP to infuse insulin in the carotid, to assess the activation of cerebral INSR *in vivo* and compare microvessel-enriched versus parenchymal fractions. We first showed that, in 6-month-old C57Bl6 mice, intracarotid administration of insulin (0.4 mg/kg or 350 nM) triggers phosphorylation of tyrosine residues (Y1150/1151) on INSRβ within 120 seconds compared to the same physiological buffer without insulin (**Fig.4D**). Faster-acting aspart insulin NovoRapid® and regular insulin Novolin®ge Toronto (350 nM) were compared and both led to the phosphorylation of INSR (**Fig. 4D and Suppl. Fig.6**). Strikingly, this activation of INSRβ was only observed in microvascular extracts, but no trace was detected in the parenchymal fraction, despite low but detectable levels of INSRs (**Fig. 4D**). This result indicates that INSR at the BBB is activated by circulating insulin and is consistent with a brain response to circulating insulin occurring predominantly at the BBB level, not in brain parenchymal cells.

To assess whether the activation of INSR is altered by Aβ and tau neuropathology, we performed the same intracarotid insulin injection experiments in 3xTg-AD mice followed by microvessel isolation and immunoblotting. While insulin induced phosphorylation of INSRβ in microvessels of 16-month-old Non-Tg mice, such activation was greatly attenuated in 3xTg-AD mice of the same age (**Fig. 4E, M**), with both types of insulin used (hexameric Novolin®ge Toronto or monomeric Aspart insulin NovoRapid®). In contrast, levels of INSRβ or pro-INSR were similar between groups (**Fig. 4F, G, M**). Replicating the data shown in **Fig. 3E** in 18-month-old 3xTg-AD mice, INSRα-B concentrations in vascular extracts were lower in this group of 15-month-old 3xTg-AD mice (**Fig. 4H, M**). The vascular INSRα-A/B ratio was higher in 3xTg-AD mice, further supporting insulin resistance at the level of the BBB (**Fig. 4I, M**).

We also assessed the effect of intracarotid insulin on other vascular proteins in NonTg and 3xTg-AD mice and no difference was found in the levels of BACE1, neprilysin, IDE and other Aβ transporters (P-gp/ABCB1, LRP1 or RAGE) (**Fig. 4J, M and Suppl. Fig.7**). However, a lower concentration of caveolin-1 was observed in 3xTg-AD compared to NonTg mice (**Fig. 4K-M**).

Results of these experiments show that AD-like neuropathology impairs the activation of vascular INSRs in the 3xTg-AD mouse model. This collection of data suggests that circulating insulin primarily interacts with INSRs exposed on the luminal side of the BBB in both human and mice and that the INSR-mediated response is blunted in AD.

### The reduction of INSR is associated with BACE1 activity in the BBB

The strong inverse correlation between BACE1 and INSR in the neurovascular compartment led us to explore whether a decrease in INSR levels could be a consequence of BACE1 protease activity, in parallel with its classical role in Aβ production (Vassar et al., 1999). The capacity of BACE1 to cleave INSR has been demonstrated in the muscle and liver, thereby contributing to insulin resistance (Meakin et al., 2018). Higher BACE1 cleavage activity of the INSR β-chain has been recently associated with cognitive impairment and T2D in a clinical cohort study (Bao et al., 2021). BACE1 levels in the cerebrovasculature are increased in AD and associated with cognitive impairment (Bourassa et al., 2019b).

To further assess BACE1 activity in microvessels, we measured the concentration of APPβ-CTF, a marker of BACE1 cleaving activity. Microvascular levels of APPβ-CTF were significantly higher in AD compared to controls based on the ABC neuropathological diagnosis (**Fig. 5A**), whereas the difference between MCI and AD based on the clinical diagnosis was not significant (p=0.053)(**Fig. 5B**). We first observed a strong positive association between vascular APPβ-CTF, INSRα-A/B ratio and BACE1 (**Fig. 5C, D**). In contrast, vascular APPβ-CTF displayed a significant inverse correlation with INSRα-B (*F*(1,40)=-10.9; p=0.003)(**Fig. 5E**) and a significant positive correlation INSRα-A/B ratio (*F*(1,40)=12.6; p=0.002). These combined data highlight the importance of BACE1 in AD in both cerebrovascular Aβ pathology and insulin resistance and suggest a role for BACE1 in cleaving vascular INSR (**Fig. 5F, G**).

**Figure 5.**
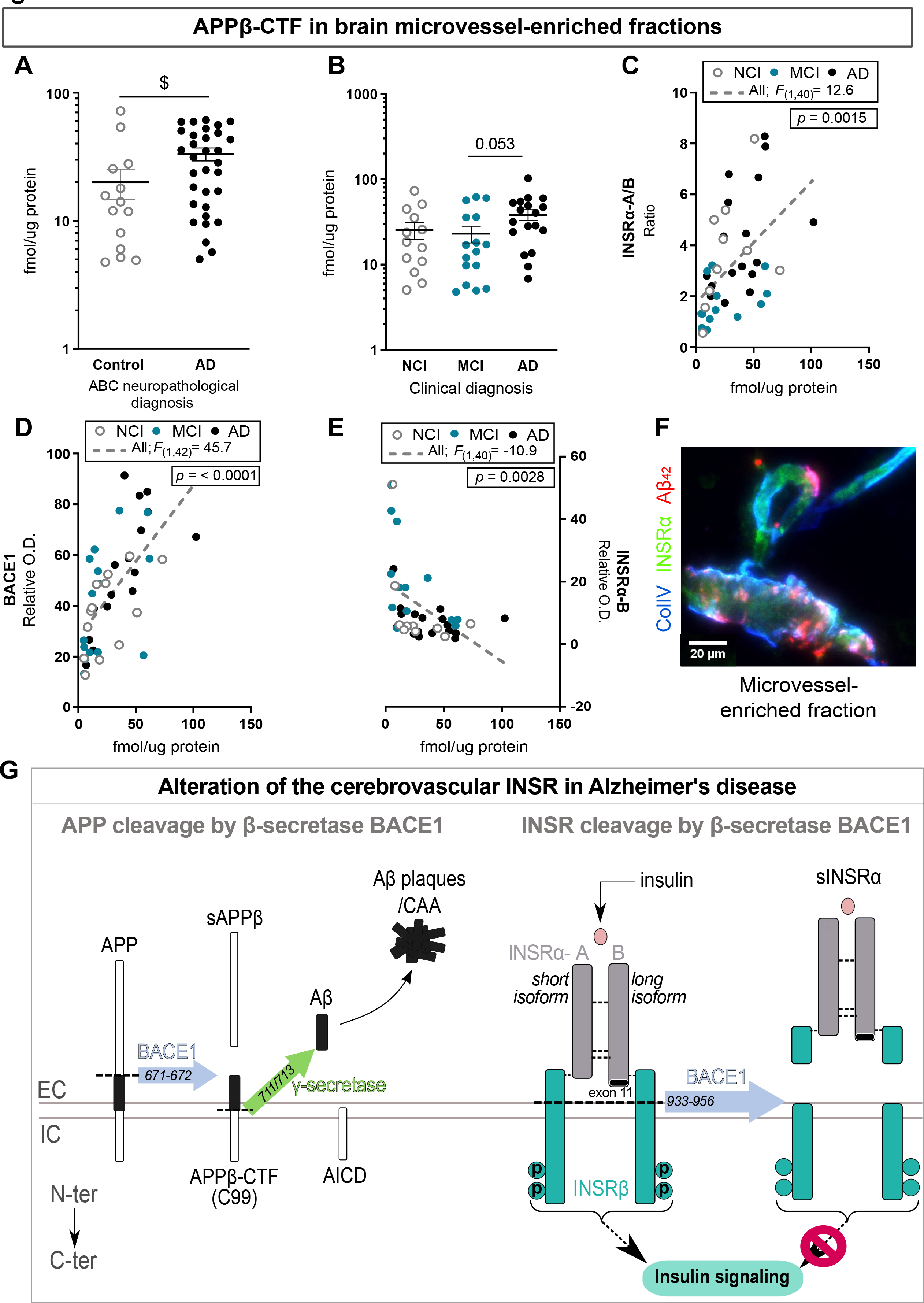
BACE1 cleavage product APPβ-CTF is negatively associated with neurovascular **levels of INSRα-B**. (A,B) Dot plots of the concentrations of APPβ-CTF in microvessels comparing participants based on neuropathological diagnosis following the ABC criteria (A) or clinical diagnosis (B). Unpaired t-test ($p < 0.05) or parametric one-way analysis of variance followed by Tukey’s post-hoc. Data were log transformed for statistical analysis and are represented as scatter plots with a logarithmic scale. Horizontal bars indicate mean ± SEM. (C-E) Correlations between the levels of vascular APPβ-CTF and BACE1, INSRαA/B ratio and INSRα-B in human brain vascular fractions. Linear regression analyses were controlled for educational level, age at death, sex and ApoE genotype. *F* ratio and p-values are shown. (F) Representative immunofluorescence labeling of human INSRα showing the colocalization with Aβ42 and collagen IV in brain microvessels of an AD patient with stage 4 cerebral amyloid angiopathy (CAA) pathology (Magnification used: 40×, scales bar: 20 μm). (G) Illustration of the role of β-secretase in cleaving APP and, hypothetically, INSR in the brain vasculature of AD patients. Abbreviations: Aβ: β-amyloid protein, ABC Dx: Neuropathological Diagnosis, AD: Alzheimer’s disease, APP: Amyloid Precursor Protein, APPβ-CTF: β-secretase-derived βAPP C-terminal fragment, AICD: Amyloid precursor protein intracellular domain, BACE1: β-site APP cleaving enzyme, CAA: Cerebral amyloid angiopathy, Clinical Dx: Clinical Diagnosis, ColIV: Collagen-IV, EC: Extracellular, IC: Intracellular, INSR: Insulin receptor, MCI: Mild cognitive impairment, NCI: Healthy controls with no cognitive impairment, O.D.: Optical density, sAPPβ: β-secretase-derived βAPP soluble fragment, sINSRα: Insulin receptor soluble fragment.

## Discussion

The present study provides evidence that INSR located at the blood-brain barrier, rather than on neurons, could underlie the brain insulin resistance observed in AD. We specifically show that (1) vascular INSR defects are linked to AD pathology and symptoms in humans and that the activation of INSR is reduced by AD neuropathology in a mouse model; (2) INSR located on the BBB is involved in the rapid response to circulating insulin, where it acts as a tyrosine kinase receptor, not a transporter. Our data suggest that pancreas-produced insulin interacts primarily with the INSR on the luminal side of the brain vasculature and that its response becomes defective in AD, possibly following cleavage by BACE1. This highlights a previously unrecognized role of INSR in the cerebral vasculature as a systemically accessible drug target in AD.

### INSR localization in brain microvessels

So far, the exact cellular localization of INSR in the brain has remained uncertain due to conflicting evidence. Autoradiography studies in rat brain sections reported prominent binding levels of ^125^I-insulin in the hippocampal formation (Dore et al., 1997), choroid plexus (Baskin et al., 1986), and hypothalamus (Baskin et al., 1983), as well as in the olfactory bulb, cerebral cortex, and cerebellum (Kar et al., 1993). A similar distribution pattern of INSRα was found using *in situ* hybridization in rat brain (Marks et al., 1990). Western blots analyses detected INSRα in the human cerebral cortex and hippocampus but not in the white matter (Frolich et al., 1998). These studies, however, did not provide insights on the cellular location of the receptor. Initial studies performed by a single group using IHC in paraformaldehyde-fixed brain sections reported a localization of INSRβ in different areas of the rat forebrain (olfactory bulb, hypothalamus, and hippocampus), largely exhibiting localization in cells resembling neurons (Moss et al., 1990; Unger et al., 1989; Unger et al., 1991). However, more recent studies with RNA sequencing in mouse and human brain cells showed that INSR is largely expressed in endothelial and glial cells (Vanlandewijck et al., 2018; Yang et al., 2021; Zhang et al., 2020). An older study with autoradiograms of ^125^I-insulin hinted toward a 130-kDa doublet corresponding to isoforms -B and -A of INSRα observed in cerebral microvessels rather than in neurons or glial cells in neonatal pigs and adult cows (Haskell et al., 1985). These discrepancies may be explained by a recurring challenge in the difficulty of finding appropriate antibodies for the INSR, and its isoforms, particularly when using human samples. This is exemplified by the number of antibodies tested here (**Suppl. Table 3**) to find a correct detection method. The present set of data indicates that most INSR isoforms in the human and mouse brain are predominantly localized in microvessels, and this was particularly the case for the isoform INSRα-B, which was virtually absent from the brain parenchyma. These results do not rule out the importance of INSR deeper in the CNS, which may interact with lower levels of insulin. Insulin-related mechanisms in the CNS might be isolated from the periphery as is the case for cholesterol and other hormones. Nonetheless, our results demonstrate that INSR are highly concentrated at the BBB and therefore likely the primary cerebral target that respond to insulin circulating in the blood after secretion by the pancreas.

### Microvascular INSR at the BBB as the site of brain insulin resistance in AD

The lower levels of vascular INSRα-B in subjects with either a neuropathological or clinical diagnosis of AD, which were inversely associated with cognitive scores, can be interpreted as a sign of brain insulin resistance in AD. Although the INSRα-B isoform was reduced, pro-INSR, INSRβ and INSRα-A did not differ significantly between groups. We found corroborating data in old 3xTg-AD mice, where INSRα-B was decreased at 18 months of age. This shift toward a higher INSRα-A/B isoform ratio at the BBB of persons with AD may have mechanistic implications. INSRα A and B isoforms can display diverging tyrosine kinase activities, as well as different affinity for insulin (Mosthaf et al., 1990; Yamaguchi et al., 1993). The B isoform of the insulin receptor has been shown to signal more efficiently than the A isoform in cultured HepG2 cells (Kellerer et al., 1992). INSR isoforms have been the subject of more scrutiny in peripheral organs (muscle, adipose tissue and liver), in cancers and in metabolic diseases, such as obesity and diabetes (Besic et al., 2015; Chettouh et al., 2013; Denley et al., 2003; Escribano et al., 2017; Kalla Singh et al., 2011; Kaminska et al., 2014; Scalia et al., 2020; Vella et al., 2018). Higher INSRα-A/B ratios are observed in adipocytes from obese patients and in the liver from T2D patients, both returning to normal after weight loss and diabetes remission (Besic et al., 2015; Kaminska et al., 2014). Therefore, a higher INSRα-A/B ratio reported in pathological conditions in insulin sensitive organs such as the liver or the muscle (Besic et al., 2015; Santoro et al., 2013) has been proposed as a molecular index of insulin resistance (Belfiore et al., 2017). In sum, the observed higher vascular INSRα-A/B ratio in AD can be interpreted as a sign of insulin resistance in the cerebrovasculature, possibly hindering the capacity of the INSR to trigger intracellular events/signaling in AD.

### Circulating insulin activates INSR in microvessels but not in the parenchyma: this activation is blunted in animal models of AD

One of the main observations reported here is that the phosphorylation of INSRβ after direct intracarotid infusion of insulin was detected at the BBB level and not in the parenchyma. Despite the presence of insulin receptors and neurons in the parenchymal fractions, no evidence of increased phosphorylation of INSRβ was detected even at a very high dose. This suggests that the physiological short-lived peak of insulin in the blood, typically observed after a meal, exerts most of its action upon INSR located on the BBB, and less in neurons. While most cells in microvessel extracts are BCEC (Bourassa et al., 2019b; Bourassa et al., 2020), which contain INSR mRNA transcripts (Vanlandewijck et al., 2018; Yang et al., 2021; Zhang et al., 2020), it is important to note that pericytes and astrocyte feet are also present in smaller amounts. It is notable that this insulin-induced INSRβ phosphorylation, observed in all control mice, was greatly attenuated in brain microvessels from 3xTg-AD mice, indicating that AD neuropathology impairs INSR signalization. This defect was observed even with the very high concentration of insulin used. Previous studies using postmortem *ex vivo* stimulation of whole brain human slices with insulin have reported lower phosphorylation of INSRβ (Y1150/1151 and Y960) in the cerebellar cortex and the hippocampal formation in AD patients (Talbot et al., 2012). However, similar studies performed in the middle frontal gyrus cortex did not find associations between p-INSRβ and diabetes, Aβ burden, tau tangle density or cognitive scores, but pS^473^ AKT1 was associated with less angiopathy (Arvanitakis et al., 2021; Arvanitakis et al., 2020). The present results suggest that INSRs located on microvessels contribute to such previously reported evidence of central insulin resistance in AD.

### Is INSR also a transporter?

Our results are not consistent with the view that INSR acts as a transporter ferrying insulin across the BBB. After insulin was shown to bind INSR at the BBB (Duffy and Pardridge, 1987; Frank and Pardridge, 1981; Frank et al., 1986; Pardridge et al., 1985), this pathway was identified as an opportunity to deliver drugs across the BBB (Pardridge et al., 1995). Evidence of transcytosis through the BBB of ^125^I-insulin or labeled monocolonal antibodies were obtained from high autoradiographic signal after systemic injection in rodents and non-human primates (Boado et al., 2018; Boado et al., 2013; Boado and Pardridge, 2017; Pardridge and Chou, 2021). Promising results were published in non-human primates, leading to clinical assays (Giugliani et al., 2018; Pardridge et al., 2018; Terstappen et al., 2021). However, this implies that the INSR is both a receptor able to trigger insulin signaling after its binding and a transporter, which is a molecularly challenging view for a tyrosine kinase receptor. Our work performed using ISCP indicates that the rate of transport through the BBB of ^125^I-insulin (0.025 ± 0.003 µl.g^-1^.s^-1^) is very low and comparable to previous estimations (0.015-0.028 ± 0.003 µl.g^-1^.s^-1^)(Banks et al., 1997; Banks et al., 2012; Rhea et al., 2018). The absence of significant antagonism following S961 co-injection is also in accordance with previous work, suggesting that INSR might not act as a transporter of insulin across the BBB (Hersom et al., 2018; Rhea et al., 2018). Therefore, while targeting the INSR may remain a viable strategy for monoclonal antibody-based BBB-targeted formulations, the present data support a scheme in which circulating insulin interact with INSRs on the BBB and triggers its known downstream signaling rather than its own transcytosis to the brain parenchyma.

### The loss of INSR is associated with a pattern of protein changes implying lower clearance and higher production of Aβ

Our report highlights the possible association between vascular INSR levels and an altered equilibrium between production and clearance of Aβ at the level of the BBB. On the one hand, INSRα-B was found to be positively associated with ABCB1 (P-gp) and LRP1, two endothelial proteins involved in Aβ efflux from the parenchyma through the BBB to the blood (Bourassa et al., 2019b; Deane et al., 2004; Storck et al., 2021; Storck et al., 2016; Zlokovic et al., 2005). Similar relationships were found with neprilysin and IDE, two key enzymes involved in Aβ degradation. Neprilysin levels were previously found to be reduced in AD microvessels (Bourassa et al., 2019b; Miners et al., 2006), while Aβ can be a substrate of IDE competing with insulin (Farris et al., 2003; Kurochkin and Goto, 1994). Published mechanistic data show that Aβ (i) can promote lysosomal degradation of INSR in 3xTg-AD mice and porcine brain capillaries (Gali et al., 2019) and (ii) act as a competitive antagonist of insulin to the INSR, thereby contributing to insulin resistance (De Felice, 2013; Xie et al., 2002). On the other hand, INSRα-B was negatively correlated with BACE1, a β- secretase initiating the cleavage of APP and formation of APPβ-CTF and Aβ (Vassar et al., 1999), both found in higher concentrations in AD brain or microvessels (Bourassa et al., 2019b; Fukumoto et al., 2002; Li et al., 2004; Stockley and O’Neill, 2007). Interestingly, using induced pluripotent stem cell (iPSC) neurons carrying APP and/or PSEN1 familial AD mutations, a recent study reports that APPβ-CTF itself may alter endosomal and intracellular trafficking, suggesting its accumulation within the neurovascular unit might contribute to BBB defects (Kwart et al., 2019). Furthermore, BACE1-generated soluble APPβ (sAPPβ) products may bind to cerebrovascular INSR and disturb it signaling (Aulston et al., 2018). Altogether, these observations put forward a scheme in which higher levels of INSRα-B in the BBB are associated with increased clearance of Aβ, lower BACE1 activity and better cognitive performance.

Other important vascular proteins such as endothelial nitric oxide synthase (eNOS or NOS3) and caveolin- 1 (Cav-1) were found to be positively associated with INSRα-B in human microvascular extracts. Cav-1 was also reduced in 3xTg-AD compared to Non-Tg mice. Previous studies showed that eNOS expression and activation in the cerebrovascular compartment are regulated by the INSR signaling cascade and by BACE1 and Aβ (Austin et al., 2013; Meakin et al., 2020; Vicent et al., 2003). Cerebral eNOS levels are reduced in AD (Jeynes and Provias, 2009). In addition, eNOS is localized in vascular endothelial cells participating, like Cav-1, in the caveola-mediated uptake of insulin, INSR trafficking and signaling in insulin-sensible organs, which are also disturbed in diabetes (Chen et al., 2019; Chen et al., 2018; Jia and Sowers, 2015; Kabayama et al., 2007; Oh et al., 2008; Wang et al., 2011). Finally, Cav-1 plays a role in INSR-mediated signaling and insulin transport in endothelial cells (Haddad et al., 2020). These patterns of association at the cerebrovascular level suggest that INSR collaborate closely with various endothelial proteins involved in the maintenance of the integrity and function of the neurovascular unit, which are also required for efficient INSR signaling.

### What is the cause of INSR decrease in AD?

Overall, these series of observations are consistent with the hypothesis that AD is accompanied by a reduction in binding sites available to circulating insulin at the level of the BBB. Several mechanisms can explain the observed decrease in INSRα-B. First, we found corroborating data in old 3xTg-AD mice, where INSRα-B was decreased at 18 months of age, with no changes in INSRβ. This suggests that age and AD neuropathology are involved. Second, it could be proposed that microvascular cells expressing INSRs are specifically lost in AD. This hypothesis is however unlikely, since we did not find a significant reduction in endothelial cells in this cohort, as assessed with claudin-5, CD31 or CypB (Bourassa et al., 2019b). A third explanation involving a change at the gene transcription level is also unlikely since the decrease was specific to the isoform INSRα-B, whereas INSRα-A, INSRβ and pro-INSR were mostly unaffected. Therefore, we postulate that post-translation modifications are more likely to underlie the observed decrease of the long isoform INSRα-B.

The strength of the correlation between vascular levels of BACE1 and INSRα-B, as well as the vascular INSRα-A/B ratio, was remarkable. A strong association was also established between INSRα-B and the concentration of APPβ-CTF, which is considered as an index of BACE1 activity (Hampel et al., 2021; Kwart et al., 2019; Vassar et al., 1999). BACE1 is probably the most studied enzyme in AD, due to its role in the production of Aβ and subsequent accumulation in the brain (Fukumoto et al., 2002; Li et al., 2004; Stockley and O’Neill, 2007). BACE1 is also expressed in microvascular endothelial cells (Devraj et al., 2016), in higher concentrations in persons with AD, correlating with cognitive impairment (Bourassa et al., 2019b). However, BACE1 is not only involved in the cleavage of APP, but it also has multiple other substrates, including INSR (Hampel et al., 2021; Koelsch, 2017). Indeed, previous reports indicate that INSR is the target of proteases, such as calpain 2 and presenilin 1 (γ-secretase), but also BACE1, which has been previously proposed as a mechanism of regulation of INSR signaling (Gaborit et al., 2021; Maesako et al., 2011; Meakin et al., 2018; Yuasa et al., 2016). Notably, studies in the liver show that BACE1 cleaves INSR between amino acids 933-956 of the β-chain, thereby reducing INSRβ levels, disturbing insulin signaling and generating a soluble fragment (sINSRα)(Gaborit et al., 2021; Meakin et al., 2018). In the liver of db/db mice, higher BACE1 protein and mRNA levels, combined with the generation of sINSRα was observed (Meakin et al., 2018). Conversely, BACE1^-/-^ mice have significantly less plasma sINSRα, higher liver INSRβ levels (Meakin et al., 2018), and improved glucose homeostasis and insulin sensitivity (Meakin et al., 2012). More recently, BACE1 cleaving activity of INSR has been related to cognitive impairment and T2D. The elevated plasma levels of BACE1 was found to correlate with the generation of sINSRα and glycemia in diabetics patients (Type I and II) (Bao et al., 2021; SolubleInsulinReceptorStudyGroup, 2007). In addition, increased plasma BACE1 activity were observed in MCI individuals (Shen et al., 2018) and higher plasma sINSRα levels can contribute to HIV-associated neurocognitive disorders (Gerena et al., 2015). Keeping these published data in mind, the series of strong correlations between BACE1 levels, BACE activity index (APPβ-CTF) and INSR levels, all measured in cerebral microvessel extracts, suggest a mechanistic scheme in which BACE1-mediated cleavage of INSR contributes to insulin resistance in AD. A favorable clinical implication is that drugs may not need to fully cross the BBB to alter INSR posttranslational modifications or to mimic the action of pancreas-secreted insulin on the brain.

## Acknowledgments

This work was supported the Canadian Institutes of Health Research (CIHR) to F.C. [grant numbers PJT 168927]. The study was supported in part by P30AG10161, P30AG72975 and R01AG15819 (D.A.B). F.C is a Fonds de recherche du Québec-Santé (FRQ-S) senior research scholar. M.L. was supported by a scholarship from the Fondation du CHU de Québec. P.B held scholarships from the Fondation du CHU de Québec, a joint scholarship from the FRQ-S and the Alzheimer Society of Canada (ASC), and a scholarship from the CIHR. V.C. was supported by a scholarship from Fonds d’Enseignement et de la Recherche (FER) from the Faculty of Pharmacy, Laval University. The authors are thankful to Gregory Klein, from the Rush Alzheimer’s Disease Research Center, for his assistance with data related to our cohort. The authors are indebted to the nuns, priests and brothers from the Catholic clergy involved in the Religious Orders Study. All procedures performed with volunteers included in this study were in accordance with the ethical standards of the institutional ethics committees and with the 1964 Helsinki Declaration. Informed consent was obtained from all individual participants included in this study.

The authors declare no competing financial interests.

## Author Contributions

M.L. performed mostly all experiments in mice and most of those in human samples, made all the analysis, figures, and tables and wrote the original draft of the manuscript. P.B. extracted the human microvessels, some mouse microvessels and made initial experiments in human samples. V.C. contributed to ISCP in mice. C.T. generated data with human samples. C.S. contributed to experiments in mice. V.E. performed a subset of immunofluorescence experiments. D.B. provided human samples and data. F.C. obtained funding, designed the experiments, and wrote the article. All authors read and approved the final manuscript.

## Abbreviations

3xTg-AD: Mouse model of genetically induced AD-like neuropathology
ABCB1 (P-gp): ATP binding cassette protein B1
AD: Alzheimer’s disease
APOE: Apolipoprotein E
APP: Amyloid precursor protein
APP-βCTF (C99): β-secretase-derived βAPP fragment
Aβ: β-amyloid peptide
BACE1: β-site APP cleaving enzyme 1
BBB: Blood-brain barrier
BCEC: Brain capillary endothelial cell
BSA: Bovine serum albumin
Cav-1: Caveolin-1
CD31 (PECAM1): Platelet endothelial cell adhesion molecule
CNS: Central nervous system;
eNOS (NOS3): Endothelial NOS or nitric oxide synthase 3
HRP: horseradish peroxidase;
GLP-1: Glucose-like peptide 1
Ht: Total homogenate;
IDE: Insulin-degrading enzyme
IHC: Immunocytochemical INSR, Insulin receptor;
ISCP: Intracarotid *in situ* cerebral perfusion;
LRP1: Low density lipoprotein receptor-related protein 1
MCI: Mild cognitive impairment;
NCI: No cognitive impairment
NeuN: Neuronal nuclei;
Non-Tg: Non-transgenic
OD: Optical density;
P: Parenchymal or microvessel-depleted fraction
PBS: Phosphate buffer saline;
RAGE: Receptor for advanced glycation endproducts;
ROS: Religious Order Study (Rush Alzheimer’s Disease Center)
RT: Room temperature;
T2D: Type-2 diabetes;
sINSRα: Insulin receptor soluble fragment
Va: Vascular or microvessel-enriched fraction

