## Supplemental Tables and Figures for "Cerebrovascular insulin receptors are defective in Alzheimerˈs disease"

### Supplementary Table 1 – Cohort Characteristics

| Characteristics | Control | AD | Statistical Analysis |
|---|---|---|---|
| *N* | 22 | 38 | ------ |
| Men, % | 41 | 29 | PC, $\chi^2$ = 0.897; p = 0.3436 |
| Mean age at death | 86.7 (4.3) | 87.5 (5.7) | MW, p = 0.5548 |
| Mean education, years | 18.3 (3.5) | 18.1 (3.1) | MW, p = 0.5236 |
| Mean MMSE | 25.0 (4.5) | 21.6 (7.9) | MW, p = 0.0791 |
| Global cognition score | -0.32 (0.81) | -0.94 (0.93)$^{\#\#}$ | MW, p = 0.0044 |
| apoE ε4 allele carriage (%) | 9 | 45$^{\&\&}$ | PC, $\chi^2$ = 8.182; p = 0.0042 |
| Clinical diagnosis NCI/MCI/AD (*n*) | 11/8/3 | 9/12/17 | ------ |
| Thal amyloid score 0/1/2/3 (n) | 7/13/2/0 | 0/3/15/20 | ------ |
| Braak score 0/1/2/3 (n) | 0/7/15/0 | 0/0/27/11 | ------ |
| CERAD score 0/1/2/3 (n) | 14/4/4/0 | 1/3/16/18 | ------ |
| Parenchymal CAA stage 0/1/2/3/4 (*n*) | 15/4/1/1/0 | 18/7/5/2/3 | PC, $\chi^2$ = 3.830; p = 0.4295 |
| Presence of chronic macroinfarcts 0/1 (*n*) | 20/2 | 33/5 | PC, $\chi^2$ = 0.224; p = 0.6363 |
| Presence of chronic microinfarcts 0/1 (*n*) | 17/5 | 34/4 | PC, $\chi^2$ = 1.627; p = 0.2021 |
| Usage of antihypertensive medications 0/1 (*n*) | 2/20 | 5/33 | PC, $\chi^2$ = 0.224; p = 0.6363 |
| Usage of diabetes medications 0/1 (*n*) | 15/7 | 33/5 | PC, $\chi^2$ = 3.032; p = 0.0816 |
| Cerebellar pH | 6.39 (0.37) | 6.29 (0.36) | MW, p = 0.2933 |
| Postmortem delay, hours | 7.93 (5.11) | 7.53 (5.15) | MW, p = 0.6652 |
| Diffuse Plaque Counts | 3.8 (8.0) | 20.3 (16.8)$^{\#\#\#\#}$ | MW, p < 0.0001 |
| Neuritic Plaque Counts | 1.3 (3.2) | 15.7 (12.5)$^{\#\#\#\#}$ | MW, p < 0.0001 |
| Neurofibrillary Tangle Counts | 0.09 (0.43) | 2.92 (8.35)$^{\#\#}$ | MW, p = 0.0096 |
| Soluble p-Tau T231/S235 | 360.7 (422.6) | 2403 (5523) | MW, p = 0.6316 |
| Soluble p-Tau S396/404 | 22.5 (13.3) | 294 (776)$^{\#}$ | MW, p = 0.0342 |
| Soluble total Tau | 803.2 (173.9) | 687.4 (212.1)$^{\$}$ | Un-W, p= 0.0138 |
| Insoluble p-Tau T231/S235 | 28.0 (94.9) | 657.2 (2503) | MW, p = 0.1537 |
| Insoluble p-Tau S396/404 | 16.4 (46.2) | 262.8 (460.7)$^{\#\#}$ | MW, p = 0.0017 |
| Insoluble Total Tau | 49.0 (18.6) | 153.8 (207.3)$^{\$\$\$}$ | Un-W, p = 0.0001 |
| Soluble $A\beta_{40}$ concentrations, fg/ug | 232.1 (595.4) | 1076 (2896)$^{\#\#\#}$ | MW, p= 0.0003 |
| Soluble $A\beta_{42}$ concentrations, fg/ug | 446.5 (527.5) | 2500 (2065)$^{\#\#\#\#}$ | MW, p < 0.0001 |
| Soluble $A\beta_{40}/A\beta_{42}$ ratio | 0.689 (0.952) | 0.667 (1.93)$^{\#\#\#}$ | MW, p= 0.0009 |
| Insoluble $A\beta_{40}$ concentrations, pg/mg | 68.8 (181.6) | 917.6 (3046)$^{\#\#\#}$ | MW, p= 0.0003 |
| Insoluble $A\beta_{42}$ concentrations, pg/mg | 426.5 (793.6) | 1577 (1128)$^{\#\#\#\#}$ | MW, p < 0.0001 |
| Insoluble $A\beta_{40}/A\beta_{42}$ ratio | 0.551 (2.06) | 0.317 (0.882) | MW, p= 0.3272 |
| Microvessel Cyclophilin B levels | 2.74 (0.77) | 2.66 (0.79) | MW, p = 0.7309 |
| Microvessel Claudin5 levels | 1.17 (0.50) | 1.16 (0.41) | MW, p = 0.6613 |
| Microvessel CD31 levels | 0.45 (0.40) | 0.41 (0.36) | MW, p = 0.7366 |

#### Supplementary Table S1 – Cohort characteristics

Participants were assigned to the "Control" or "AD" group based on ABC scores in the parietal cortex (see Method). Unless noted, all measures are from the parietal cortex, except the brain pH which was measured in the cerebellum. Aβ peptide concentrations were determined by ELISA and Tau by Western blot. Values are expressed as means (SD) unless specified otherwise. Statistical analysis (compared to controls): Unpaired t-test with Welch correction, $^{\$}p < 0.05$, $^{\$\$\$}p < 0.001$, Mann–Whitney test, $^{\#}p < 0.05$, $^{\#\#}p < 0.01$, $^{\#\#\#}p < 0.001$, $^{\#\#\#\#}p < 0.0001$, Pearson's χ2 contingency test, $^{\&\&}p < 0.01$. Claudin-5 and CD31 data in microvascular extracts were normalized with Cyclophilin B as loading control.

Abbreviations: Aβ: β-amyloid peptides, AD: Alzheimer's disease, ApoE: Apolipoprotein E genotype, CAA: Cerebral amyloid angiopathy, CD31: Platelet endothelial cell adhesion molecule; CERAD: Consortium to Establish a Registry for Alzheimer's Disease, MCI: Mild cognitive impairment, MMSE: Mini-Mental State Examination, MW: Mann-Whitney test, NCI: Healthy controls with no cognitive impairment, PC: Pearson's χ2 contingency test, Un-W: Unpaired t-test with Welch correction.

### Supplementary Table 2 - List of antibodies used for Western blotting and immunofluorescence

| Protein | Role/localization | Host | Dilution | | Company | Catalog number |
|---|---|---|---|---|---|---|
| | | | WB | IF | | |
| **ABCB1(P-gp)** | BBB transporter | Rabbit | 1:2000-1:5000 | | Abcam | 170904 |
| **Aβ$_{40}$** | Aβ peptide | Mouse | | 1:100 | BioLegend | 805401 |
| **Aβ$_{42}$** | | Mouse | | 1:100 | BioLegend | 805501 |
| **β-actin** | Cytoskeletal ubiquitously expressed protein | Mouse | 1:10000 | | Applied Biological Materials | G043 |
| **Akt 1/2/3** | Protein kinase B/Akt | Rabbit | 1:1000 | | Santa Cruz Biotechnology | 8312 |
| **Akt1 (D9R8K)** | | Rabbit | 1:1500 | | Cell signaling | 75692 |
| **Akt2 (D6G4)** | | Rabbit | 1:1250 | | Cell signaling | 3063 |
| **p-Akt1 (Thr308)** | | Rabbit | 1:1500 | | Cell signaling | 9275 |
| **p-Akt1(2-3) Ser473** | | Rabbit | 1:1000 | | Cell signaling | 9271 |
| **BACE1** | β-secretase contributing to the production of Aβ | Rabbit | 1:500-1:1000 | | Abcam | 108394 |
| **Caveolin-1** | Caveolae plasma membranes | Rabbit | 1:1000 | | Cell signaling | 3267 |
| **CD31** | EC intercellular junctions | Rabbit | 1:1000-:2000 | | Abcam | 28364 |
| **Claudin-5** | | Rabbit | 1:2000-1:5000 | 1:100 | Santa Cruz Biotechnology | 28670; discontinued |
| **Collagen IV** | Basal lamina | Goat | | 1:500 | Sigma-Aldrich | AB769 |
| | | Rabbit | | 1:500 | Abcam | ab6586 |
| **Cyclophilin B** | Ubiquitous cellular proteins | Rabbit | 1:1000 | | Abcam | 16045 |
| **eNOS** | Endothelial nitric oxide synthase (NOS3) | Mouse | 1:1000 | | Biosource | 610297 |
| **GAP43** | Schwann cells | Rabbit | 1:1000 | | Abcam | 75810 |
| **GSK3β** | Glycogen Synthase Kinase-3 β | mouse | 1:1000 | | Biosource | 610202 |
| **p-GSK3β (ser9)** | | Rabbit | 1:1000 | | Cell signaling | 9336 |
| **p-GSK3β (Tyr216)** | | Rabbit | 1:1000 | | Abcam | 75745 |
| **IDE** | Insulin degrading enzyme | Rabbit | 1:1000 | | Abcam | 133561 |
| **INSRα** | Human INSR extracellular α chain between His28-Lys944 | Goat | 1:500-1:1000 | 1:100-1:200 | R&D Systems | AF1544 |
| **INSRβ (E9L5V)** | INSR cytosolic β chain and pro-INSR | Rabbit | 1:500 | 1:200 | Cell signaling | 23413 |
| **p-IGF1Rβ/INSRβ (19H7)** | Phosphorylated IGF1Rβ at Tyr1135/1136 or INSRβ at Tyr1150/1151 | Rabbit | 1:500-1:1000 | | Cell signaling | 3024 |
| **LRP1** | BBB receptor | Rabbit | 1:1250 | | Abcam | 92544 |
| **MAPK p44/p42 (Erk 1/2)** | Mitogen-activated protein kinases signaling pathway | Rabbit | 1:1000 | | Cell Signaling | 9102S |
| **p-MAPK p44/p42 (Thr202/Ty204)** | | Rabbit | 1:1500 | | Cell Signaling | 9101S |
| **Neprilysin** | Aβ-degrading enzyme | Rabbit | 1:2000 | | Abcam | 79423 |
| **NeuN** | Neuronal nuclei | Rabbit | 1:1000 | 1:500 | Abcam | 177487 |
| **RAGE** | BBB receptor | Rat | 1:2000 | | R&D Systems | MAB11795 |
| **Total soluble Tau (Tau13)** | Tau cytoskeletal protein | Mouse | 1:1000 | | Covance/Biolegend | 835201 |
| **Total insoluble Tau (Tau640-680)** | | Rabbit | 1:3000 | | Osenses | OST00329W |
| **p-Tau T231/S235 (AT180)** | | Mouse | 1:1000 | | Thermo Fisher Scientific | MN1040 |
| **p-Tau S396/404 (PHF1)** | | Mouse | 1:250 | | Davies | |
| **mTor** | Mammalian target of rapamycin | Rabbit | 1:1000 | | Cell Signaling | 2983 |
| **p-mTor (D9C2) (Ser2448)** | | Rabbit | 1:1000 | | Cell Signaling | 5536S |
| **β-Tubulin III** | Neuronal microtubules | Rabbit | 1:2000 | | Sigma-Aldrich | T2200 |
| **Secondary antibodies** | | | | | | |
| **anti-Rabbit-HRP** | HRP-conjugated | Goat | 1:20 000-1:60 000 | | Jackson | 111-035-144 |
| **anti-Mouse-HRP** | | Goat | 1:20 000-1:60 000 | | Jackson | 115-035-166 |
| **anti-Goat-HRP** | | Donkey | 1:30 000-1:40 000 | | Jackson | 705-035-147 |
| **anti-Goat-AF555** | Alexa Fluor-conjugated | Donkey | | 1:500 | Thermo Fisher Scientific | A21432 |
| **anti-Goat-AF647** | | Donkey | | 1:500 | Thermo Fisher Scientific | A32849 |
| **anti-Goat-AF790** | | Donkey | | 1:500 | Jackson | 705-655-147 |
| **anti-Mouse-AF555** | | Donkey | | 1:500 | Thermo Fisher Scientific | A31570 |
| **anti-Mouse-AF647** | | Donkey | | 1:500 | Thermo Fisher Scientific | A31571 |
| **anti-Rabbit-AF555** | | Donkey | | 1:500 | Thermo Fisher Scientific | A31572 |
| **anti-Rabbit-AF790** | | Donkey | | 1:500 | Thermo Fisher Scientific | A11374 |

**Supplementary Table S2 – List of antibodies used in this study for Western blotting and immunofluorescence**

Abbreviations: Aβ: β-amyloid peptides, ABCB1(P-gp), AF: Alexa Fluor, ATP Binding Cassette Subfamily B Member 1/P-glycoprotein, Akt: Protein kinase B, BACE1: β-secretase, BBB: Blood–brain barrier, CD31: Platelet endothelial cell adhesion molecule, EC: Endothelial cell, eNOS: Endothelial nitric oxide synthase, GAP43: Growth Associated Protein 43, GSK3β: Glycogen synthase kinase 3 β, HRP: horseradish peroxidase, IDE, Insulin-degrading enzyme, IF: Immunofluorescence, INSR: Insulin receptor, LRP1: Low-density lipoprotein receptor-related protein 1, MAPK: Mitogen-activated protein kinase, mTor: Mammalian target of rapamycin, NeuN: Neuronal nuclei, RAGE: Receptor for advanced glycation end products, WB: Western blot.

### Supplementary Table 3 - List of INSR and IRS-1 antibodies tested in microvascular extracts

| Proteins | Host | Size (kDa) | Mouse | Human | Company | Catalog number |
|---|---|---|---|---|---|---|
| **p-IGF1Rβ/INSRβ (19H7)**<br>(phosphorylated IGF1Rβ at Tyr1135/1136 or INSRβ at Tyr1150/1151) | Rabbit | 95 | ++ | Not applicable | Cell signaling | 3024 |
| **p-INSRβ (10C3)**<br>(phosphorylated INSRβ at Tyr1150/1151) | Mouse | 95 | + | Not applicable | Santa Cruz Bio. | 81500 |
| **p-IGF1Rβ/INSRβ**<br>(phosphorylated IGF1Rβ at Tyr1131/1135/1136 or INSRβ at Tyr1146/1150/1151) | Rabbit | 95 | - | Not applicable | Genetex | GTX25681 |
| **p-IGF1Rβ/INSRβ**<br>(phosphorylated IGF1Rβ at Tyr1131 or INSRβ at Tyr1146) | Rabbit | 95 | - | Not applicable | Cell signaling | 3021 |
| **p-IGF1Rβ/INSRβ**<br>(phosphorylated IGF1Rβ or INSRβ at Tyr1158/1162/1163) | Rabbit | 95 | - | Not applicable | Milipore | 07-841 |
| **p-INSRβ**<br>(phosphorylated INSRβ at Tyr1160, no indication with IGF1R cross reaction) | Rabbit | 95 | - | Not applicable | Phosphosolution | p168-1160 |
| **p-INSRβ (14A4)**<br>(phosphorylated INSRβ at Tyr1345) | Rabbit | 95 | - | Not applicable | Cell signaling | 3026 |
| **INSRβ (E9L5V)**<br>(INSR cytosolic β chain and pro-INSR, no indication with IGF1R cross reaction) | Rabbit | 95 and 210 | +++ | ++ | Cell signaling | 23413 |
| **INSRβ (L55B10)**<br>(INSR cytosolic β chain at Tyr999, not supposed to cross react with IGF1R) | Mouse | 95 | + | - | Cell signaling | 3020 |
| **INSRβ (4B8)**<br>(INSR cytosolic β chain at Tyr999, not supposed to cross react with IGF1R) | Rabbit | 95 | ++ | - | Cell signaling | 3025 |
| **INSRβ (C19)**<br>(INSR cytosolic β chain) | Rabbit | 95 | ++ | - | Santa Cruz Bio. | 711 discontinued |
| **INSRβ (CT-3)**<br>(INSR cytosolic β chain, not supposed to cross react with IGF1R) | Mouse | 95 | ++ | + with IgG1 specific secondary antibodies only | Santa Cruz Bio. | 57342 |
| **INSRβ**<br>(INSR transmembrane and cytosolic domains amino acids 939-1347, no indication with IGF1R cross reaction) | Rabbit | 95 | +<br>very weak | - | Thermo Fisher | PA5-27334 |
| **INSRα**<br>(human INSR extracellular α chain between His28-Lys944) | Goat | 135 | ++ | ++ | R&D Systems | AF1544 |
| **INSRα**<br>(extracellular N-terminal domain amino acids 28-57) | Rabbit | 140 | - | - | Abgent | AR7653a |
| **INSRα (83-14)**<br>(human INSR extracellular α chain between His28-Lys944) | Mouse | 135 | - | - | Thermo Fisher | AHR0221 |
| **INSRα (83-22)**<br>(human INSR extracellular α chain) | Mouse | 150 | - | - | BioXcell | BE0338 |
| **INSR**<br>blocking peptide (33R-2396) | Rabbit | 155 | - | - | Fitzgerald | 70R-8501 |
| **p-IRS1 (Ser612)** | Rabbit | 180 | - | - | Cell signaling | 2386 |
| **p-IRS1 (Tyr632)** | Rabbit | 180 | +/-<br>2 bands at 130/200 kDa | - | Abcam | 109543 |
| **p-IRS1 (Ser636/639)** | Rabbit | 180 | - | - | Cell signaling | 2388 |
| **p-IRS1(Ser1101)** | Rabbit | 180 | +/-<br>very weak | - | Cell signaling | 2385 |
| **IRS1** | Rabbit | 180 | - | - | Cell signaling | 2382 |
| **IRS1 (E-12)** | Mouse | 180 | - | non specific band at 50 kDa | Santa Cruz Bio. | 8038 |

#### Supplementary Table 3 – List of INSR and IRS-1 antibodies tested in microvascular extracts

Abbreviations: INSR: Insulin receptor, IRS: Insulin receptor substrate.

**Supplementary Figure 1 - Schematic summary of the intracarotid in situ cerebral perfusion (ISCP) followed by the capillary extraction technique**

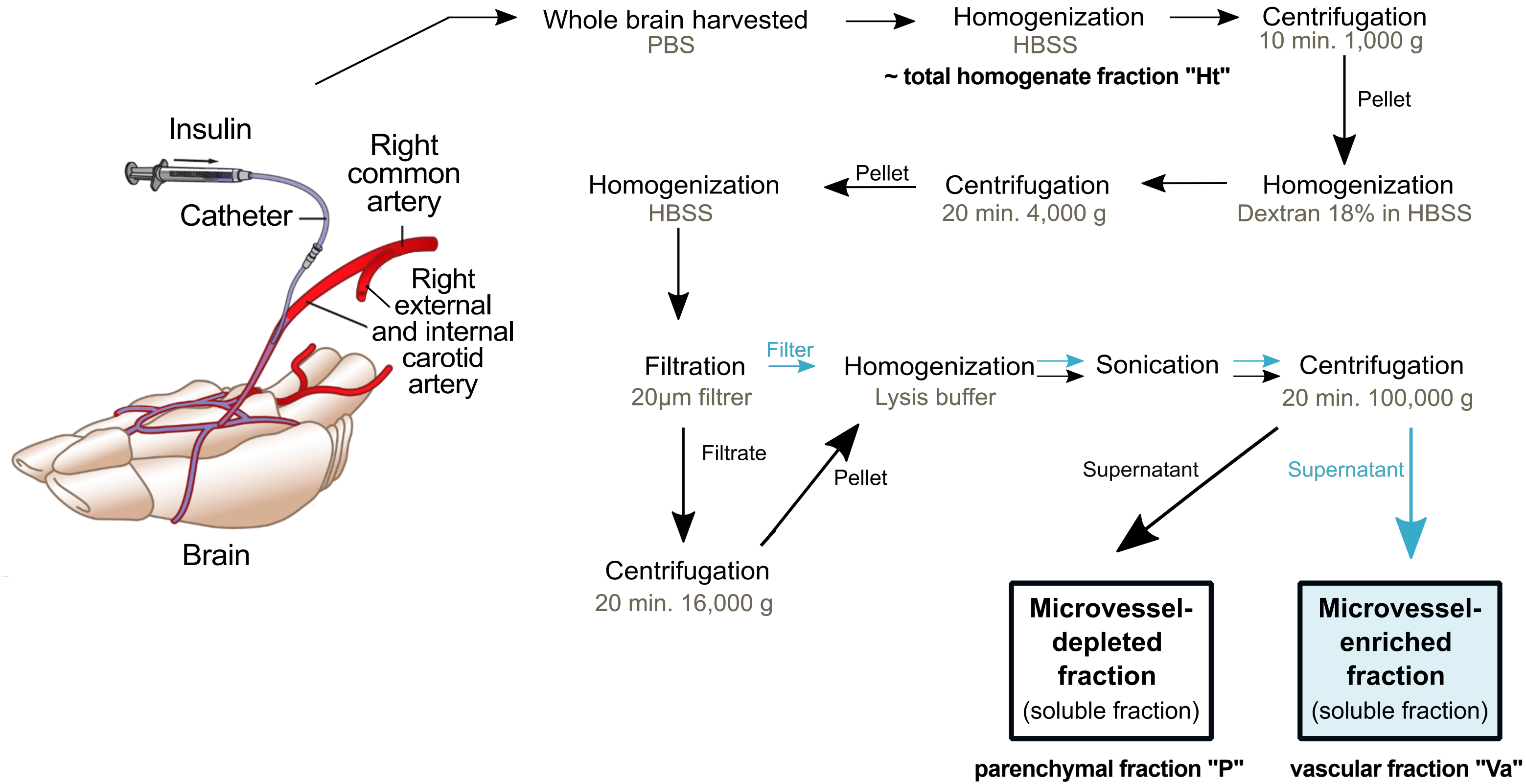

**Supplementary Figure S1 – Schematic summary of the intracarotid *in situ* cerebral perfusion (ISCP) followed by the capillary extraction technique**

Intracarotid *in situ* cerebral perfusion followed by capillary extraction protocol illustration. Workflow leading to the separation of the microvessel-enriched and the microvessel-depleted parenchymal fractions, respectively illustrated in the blue box and the white box, from whole homogenates of cerebral cortex.

Abbreviations*:* HBSS: Hanks' balanced salt solution, Ht: Total Homogenate, P: Microvessel-depleted parenchymal fraction, PBS: Phosphate buffer saline, Va: Vascular fraction enriched in microvessels.

**Supplementary Figure 2 - Western blots in human brain microvascular extracts**

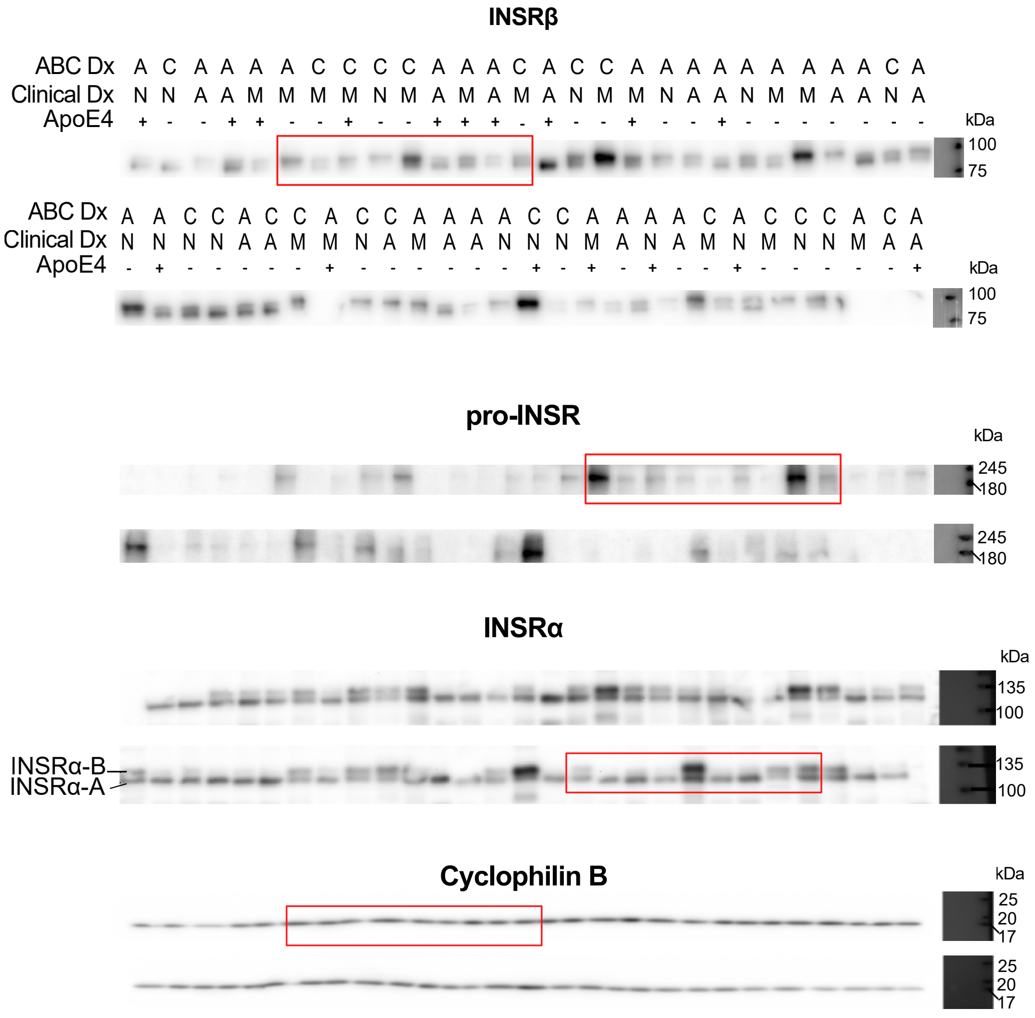

#### Supplementary Figure S2 – Western blots in human brain microvascular extracts

Uncropped gels of immunoblotting experiments in human brain microvessels. The clinical and neuropathological diagnoses are given above each sample. An equal amount (8 µg) of proteins per sample was loaded. Red rectangles indicate the bands that were taken as representative photo examples in Fig.1.

Abbreviations: A: Alzheimer's Disease, ABC Dx: ABC neuropathological diagnosis, ApoE4, Apolipoprotein E4 genotype, C: Control, Clinical Dx: Clinical diagnosis, INSR: Insulin receptor, M: mild cognitive impairment, N: Healthy control with no cognitive impairment.

**Supplementary Figure 3 - INSRα-B is significantly decreased in microvessels in individuals classified with higher ABC subscores (Thal, Braak and CERAD) and cerebral amyloid angiopathy (CAA) stages**

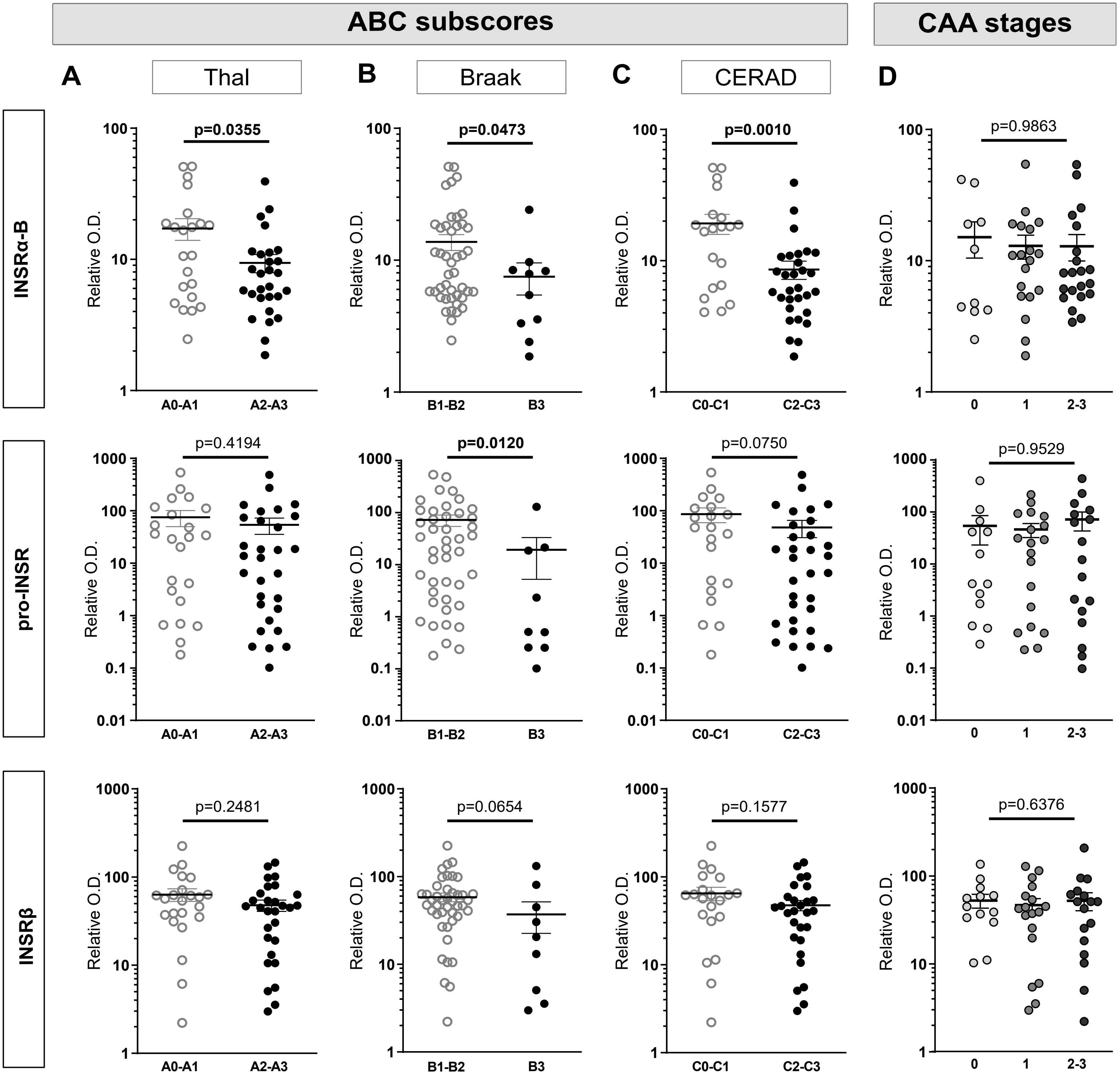

**Supplementary Figure S3 – INSRα-B is significantly decreased in microvessels in individuals classified with higher ABC subscores (Thal, Braak and CERAD) and cerebral amyloid angiopathy (CAA) stages.**

(A-D) Dot plots of the concentrations of INSR isoforms in microvessels comparing participants based on their neuropathological diagnosis following the ABC criteria and classified CAA stages for INSRα-B, pro-INSR, and INSRβ levels. "A"-Thal score assessing phases of Aβ plaque accumulation (A), "B"-Braak score assessing neurofibrillary tangle pathology (B) and "C"-CERAD score assessing neuritic plaque pathology (C). Parenchymal CAA stages in the parietal cortex were determined in the angular gyrus (D).

Unpaired t-test, Mann-Whitney or parametric one-way analysis of variance. Data were log transformed for statistical analysis and are represented as scatter plots with a logarithmic scale. Horizontal bars indicate mean ± SEM.

Abbreviations: CAA: Cerebral amyloid angiopathy, INSR: Insulin receptor, O.D.: Optical density.

### Supplementary Figure 4 - Western blots of INSRα in human brain microvessel-depleted fractions (brain parenchyma)

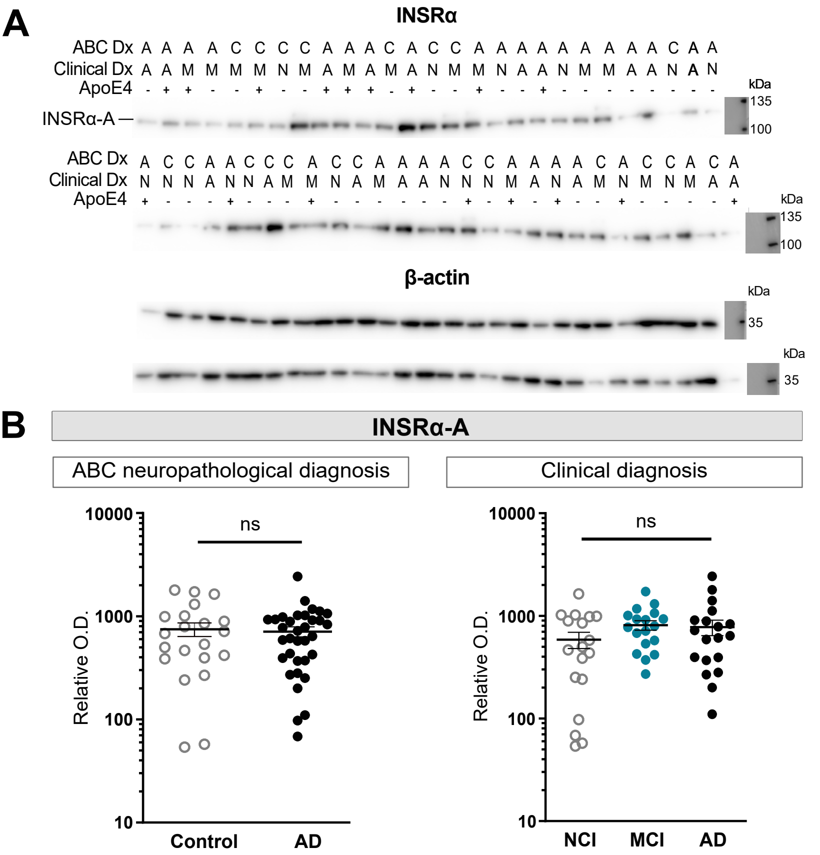

**Supplementary Figure S4 – Western blots of INSRα in human brain microvessel-depleted fractions (brain parenchyma)**

(A) Western blots of INSRα showing only INSRα-A in parenchymal fraction from the parietal cortex along with β-actin. (B) Dot plots of the concentrations of INSRα-A in microvessel-depleted fractions comparing participants based on the neuropathological diagnosis following the ABC criteria or clinical diagnosis. Unpaired t-test with Welch correction or Welch's ANOVA (ns, not significant). Data were log transformed for statistical analysis and are represented as scatter plots with a logarithmic scale. Horizontal bars indicate mean ± SEM.

Abbreviations: A-AD: Alzheimer's Disease, ABC Dx: ABC neuropathological diagnosis, ApoE: Apolipoprotein E genotype, C: Control, Clinical Dx: Clinical diagnosis, INSR: Insulin receptor, M-MCI: Mild cognitive impairment, N-NCI: Healthy control with no cognitive impairment, O.D.: Optical density.

**Supplementary Figure 5 - Linear regressions between INSR in brain microvessels and antemortem cognitive evaluation**

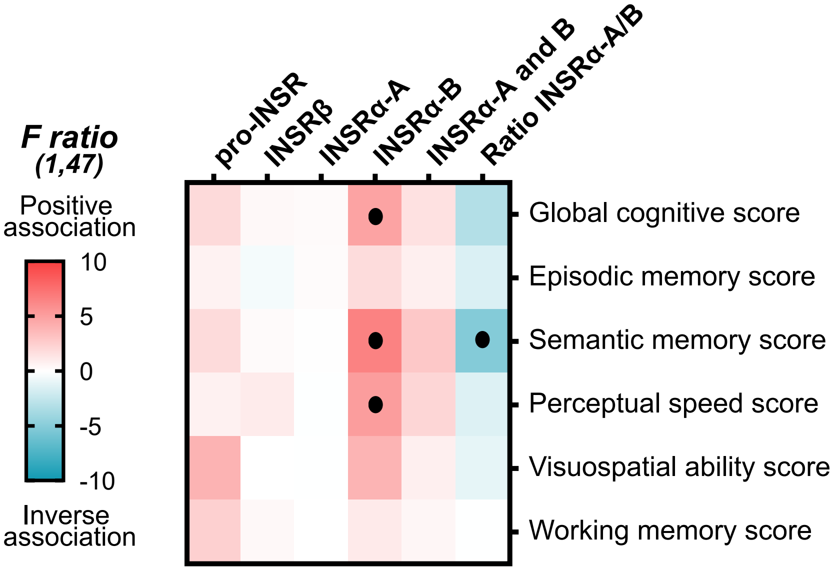

**Supplementary Figure S5 – Linear regressions between INSR in brain microvessels and antemortem cognitive evaluation**

Correlations between cerebrovascular INSRs and *antemortem* cognitive. Linear regressions analyses were controlled for educational level, age at death, sex and ApoE genotype, and were performed to generate *F ratios* and p-values in the heatmap (•$p < 0.05$). Red and blue highlighted cells respectively indicate significant positive and negative correlations.

Abbreviations: INSR: Insulin receptor.

### Supplementary Figure 6 - INSR phosphorylation after *in situ* cerebral perfusion (ISCP) of insulin

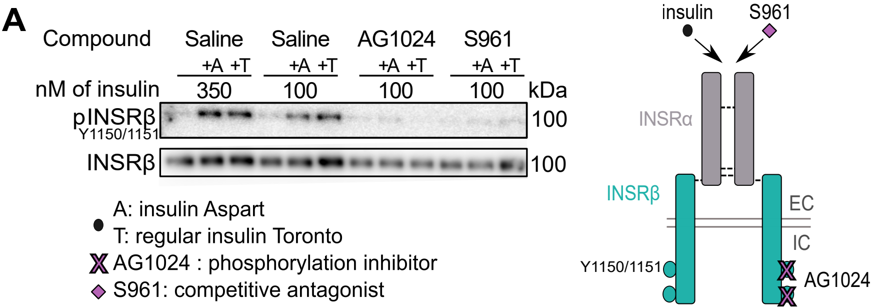

**Supplementary Figure S6 – INSR phosphorylation after *in situ* cerebral perfusion (ISCP) of insulin.**

(A) Illustration and representative Western blots of experiments showing phosphorylated-INSRβ in vascular fractions following insulin perfusion (100 nM and 350 nM) by ISCP in 6-month-old C57Bl6 mice and blunted with coperfused INSRβ phosphorylation inhibitor (AG1024) or INSR antagonist (S961) (n=2-4).

Abbreviations: A: insulin Aspart, EC: extracellular, IC: intracellular, INSR: Insulin receptor, T: regular insulin Toronto.

#### Supplementary Figure 7 - Microvessels proteins after *in situ* cerebral perfusion of insulin

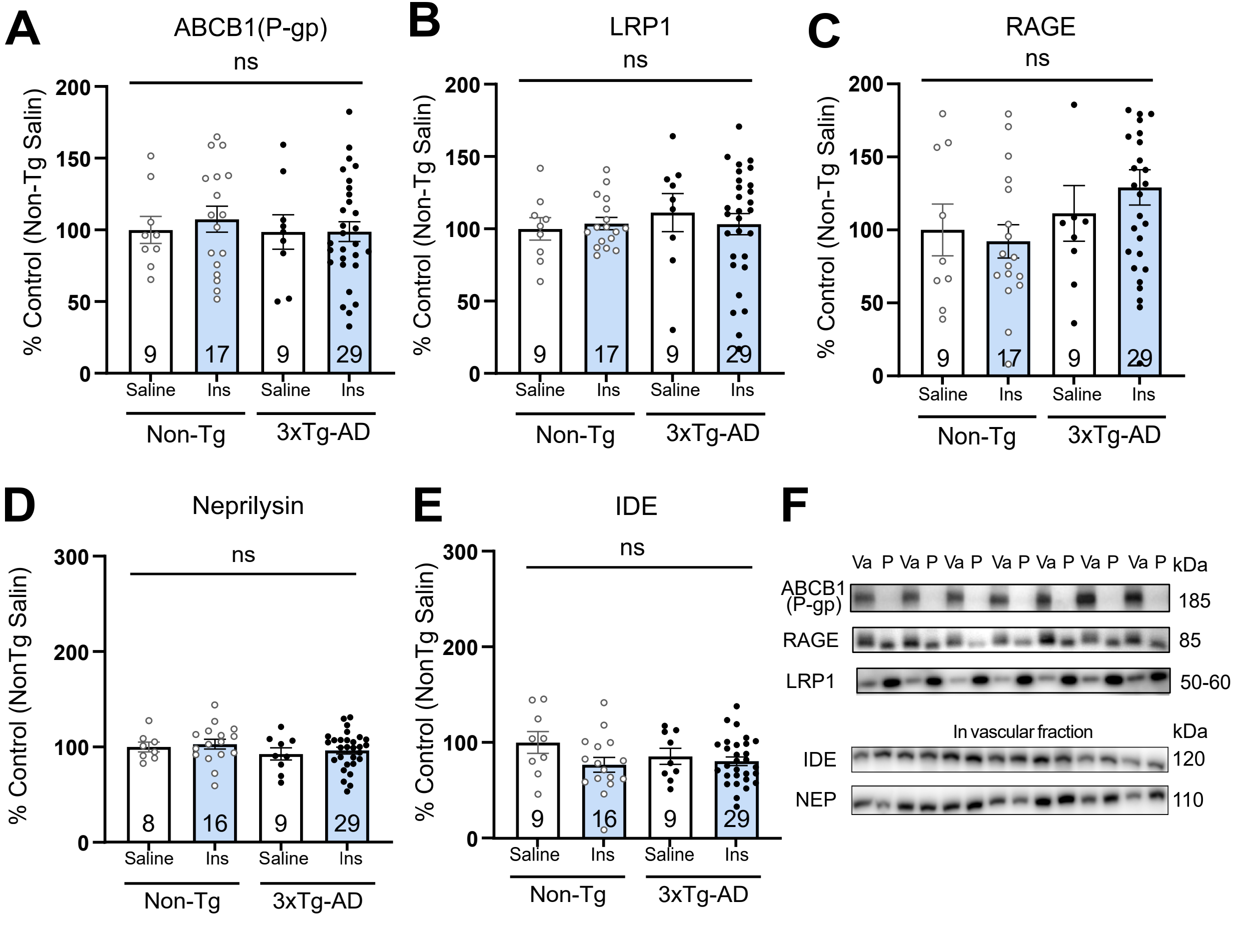

**Supplementary Figure S7 – Microvessel proteins after *in situ* cerebral perfusion (ISCP) of insulin.**

(A-F) Dot plots of the concentrations of cerebrovascular proteins in microvessels comparing mice based on genotype and insulin perfusion. One-way or two-way analysis of variance were performed (ns, non-significant). Data were log transformed for statistical analysis. Horizontal bars indicate mean ± SEM. Representative Western blots of consecutive bands are shown from the samples in different randomized gels (F).

Abbreviations: 3xTg-AD: tri-transgenic mice, ABCB1(P-gp): ATP Binding Cassette Subfamily B Member 1/P-glycoprotein, IDE: Insulin degrading enzyme, Ins: Insulin perfusion, INSR: Insulin receptor, LRP1: Low density lipoprotein receptor-related protein 1, NEP: Neprilysin, Non-Tg: Non-transgenic mice, O.D.: Optical density, P: Microvessel-depleted parenchymal fraction, RAGE: Receptor for Advanced Glycation End products, Va: Vascular fraction enriched in microvessels.
